# Identification of genes conferring individual-level variation responsible for metabolic dysfunction-associated steatohepatitis using single-cell eQTL analysis

**DOI:** 10.1101/2024.09.20.614203

**Authors:** Sung Eun Hong, Seon Ju Mun, Young Joo Lee, Taekyeong Yoo, Kyung-Suk Suh, Keon Wook Kang, Myung Jin Son, Won Kim, Murim Choi

**Affiliations:** Department of Biomedical Sciences, Seoul National University College of Medicine, Seoul 03080, Republic of Korea; Stem Cell Convergence Research Center, Korea Research Institute of Bioscience and Biotechnology (KRIBB), Daejon 34141, Republic of Korea; College of Pharmacy and Research Institute of Pharmaceutical Sciences, Seoul National University, Seoul 08826, Republic of Korea; Department of Surgery, Seoul National University College of Medicine, Seoul 03080, Republic of Korea; Department of Functional Genomics, Korea University of Science & Technology (UST), Daejeon 34113, Republic of Korea; Division of Gastroenterology and Hepatology, Department of Internal medicine, Seoul Metropolitan Government Boramae Medical Center, Seoul National University College of Medicine,, Seoul 07061, Republic of Korea

## Abstract

Metabolic dysfunction-associated steatotic liver disease (MASLD) is increasingly recognized for its medical and socioeconomic impacts, driven by diverse genetic and environmental factors. To address the urgent need for individually tailored therapies, we performed single-cell expression quantitative trait loci (sc-eQTL) analysis on liver biopsies from 25 MASLD patients and 23 controls. This approach identified over 3,500 sc-eQTLs across major liver cell types and cell state-interacting eQTLs (ieQTLs) with significant enrichment for disease heritability (for MASLD trait, ieQTL enrichment odds ratio = 10.27). We integrated transcription factors (TFs) as upstream regulators of ieQTLs, revealing 601 functional units (“quartets”) composed of TFs, cell states, ieSNPs, and ieGenes. From these results, we pinpoint the loss of an eQTL in *EFHD1* during hepatocyte maladaptation associated with genotype-specific regulation by FOXO1, further contributing to the risk of MASLD. Our approach underscores the role of eQTL analysis in capturing crucial genetic variations that influence gene expression and clinical outcomes in complex diseases.

## Introduction

As a manifestation of metabolic dysregulation, metabolic dysfunction-associated steatotic liver disease (MASLD) affects more than a quarter of the adult population worldwide, and thereby posing a substantial burden of end-stage liver disease^1^. MASLD collectively represents a spectrum of liver conditions ranging from isolated steatosis to metabolic dysfunction-associated steatohepatitis (MASH), with progression to cirrhosis and hepatocellular carcinoma in a subset of patients^1,2^. This high degree of inter-individual variability in disease progression hampers the provision of optimal, patient-specific treatment options; furthermore, the molecular genetic mechanisms underlying the variability are not fully understood^3^.

Susceptibility to and severity of MASLD are known to be influenced by the interplay between genetic and environmental risk factors^1,4^. The estimated heritability ranges from 20% to 50%^5^, and genome-wide association studies (GWAS) and rare variant analyses have identified a number of genetic variants showing significant associations^3,6–11^. The top variants in *PNPLA3* and *HSD17B13* are protein-altering, rendering them attractive targets for novel precision medicine drugs^12–14^. However, most variants are located outside the coding region, hindering their biological interpretation and clinical translation^15^.

Non-coding GWAS variants cluster around regulatory elements and play an important role in regulating molecular phenotypes such as gene expression^16^. Mapping of expression quantitative trait loci (eQTL) identifies genes regulated by genetic variants, thus providing a mechanistic link between genetic variants and phenotypes^17^. Large-scale eQTL studies^18,19^ have profiled the genetic architecture of gene expression in healthy, adult, and steady-state tissues; however, such conventional eQTLs do not fully recapitulate complex and subtle differences in physiological status, and cannot reveal regulatory variation specific to disease contexts^20,21^. Previous attempts to discover eQTLs in a context-specific manner, such as in relation to development^22–24^, immune stimuli^25^, drug treatment^26^, cell stress^27^, or disease^28,29^, have successfully identified novel disease-associated regulatory variants and their target genes^29^. Our previous study identified eQTLs specific to MASLD donors (*i.e.*, response-eQTLs) and suggested *AGXT2* as a potential genotype-dependent therapeutic target^29,30^. Despite these successes, dynamic eQTL discovery via bulk-based methods remains hindered by the difficulty in acquiring disease-associated samples and challenges in modeling continuous contexts^20^.

Single-cell RNA-sequencing (scRNA-seq) can simultaneously and unbiasedly quantify gene expression and cell states^31–33^, allowing novel dynamic eQTL discovery^34^. Therefore, single-cell eQTL (sc-eQTL) analyses of blood^35–37^, brain^38,39^ or induced pluripotent stem cells^40,41^ are able to reveal context-specific eQTLs associated with various diseases. Many sc-eQTL studies have employed statistical approaches similar to bulk-eQTLs, or linear regression on pseudobulk expression aggregated from discrete clusters. Recently, an approach has been proposed that actively incorporates the dynamic cell state information obtained from single-cell-level data to test for interactions with the regulatory effects of genetic variants (*i.e.,* cell state-interacting eQTLs; ieQTLs)^37^, which we have applied in this study.

Here we performed single-nucleus RNA-seq (snRNA-seq) on liver tissues biopsied from patients with various stages of MASLD. This enabled us to characterize cells from various contexts and states. We adopted the Poisson mixed-effects (PME) model to call conventional eQTLs and ieQTLs at single-cell resolution, then further analyzed the ieQTLs for their disease relevance and identified target genes with therapeutic potential. Our study illuminates the importance and utility of the sc-eQTL approach in providing genetic insights into pathogenic mechanisms, such as in liver disease, and suggesting novel causal genes for genotype-dependent treatment strategies.

## Results

### snRNA-seq reveals transcriptomic alterations within and across cell types in MASLD

We performed 10x-based single-nucleus RNA-sequencing (snRNA-seq) on liver tissues obtained from 30 MASLD patients and 24 non-MASLD individuals. This sample size is based on a power estimation suggesting 82.9% of power to detect variants with an effect size of at least 0.8 could be achieved using 50 samples^27^ (Supplementary Methods and Supplementary Fig. 1). To account for the wide clinical spectrum of the disease, the study participants covered diverse stages ranging from no MASLD or isolated steatosis (MASL) through MASH with mild fibrosis (early MASH; eMASH) to MASH with advanced fibrosis (advanced MASH; aMASH) (Fig. 1a, Supplementary Methods and Supplementary Table 1). Following stringent quality control steps^36,37,42^, 249,233 nuclei from 48 donors were retrieved for calling conventional eQTLs (liver- eQTLs) and cell-state interacting eQTLs (ieQTLs) (Fig. 1a, Supplementary Fig. 2, Supplementary Tables 1 and 2).

**Figure 1.**
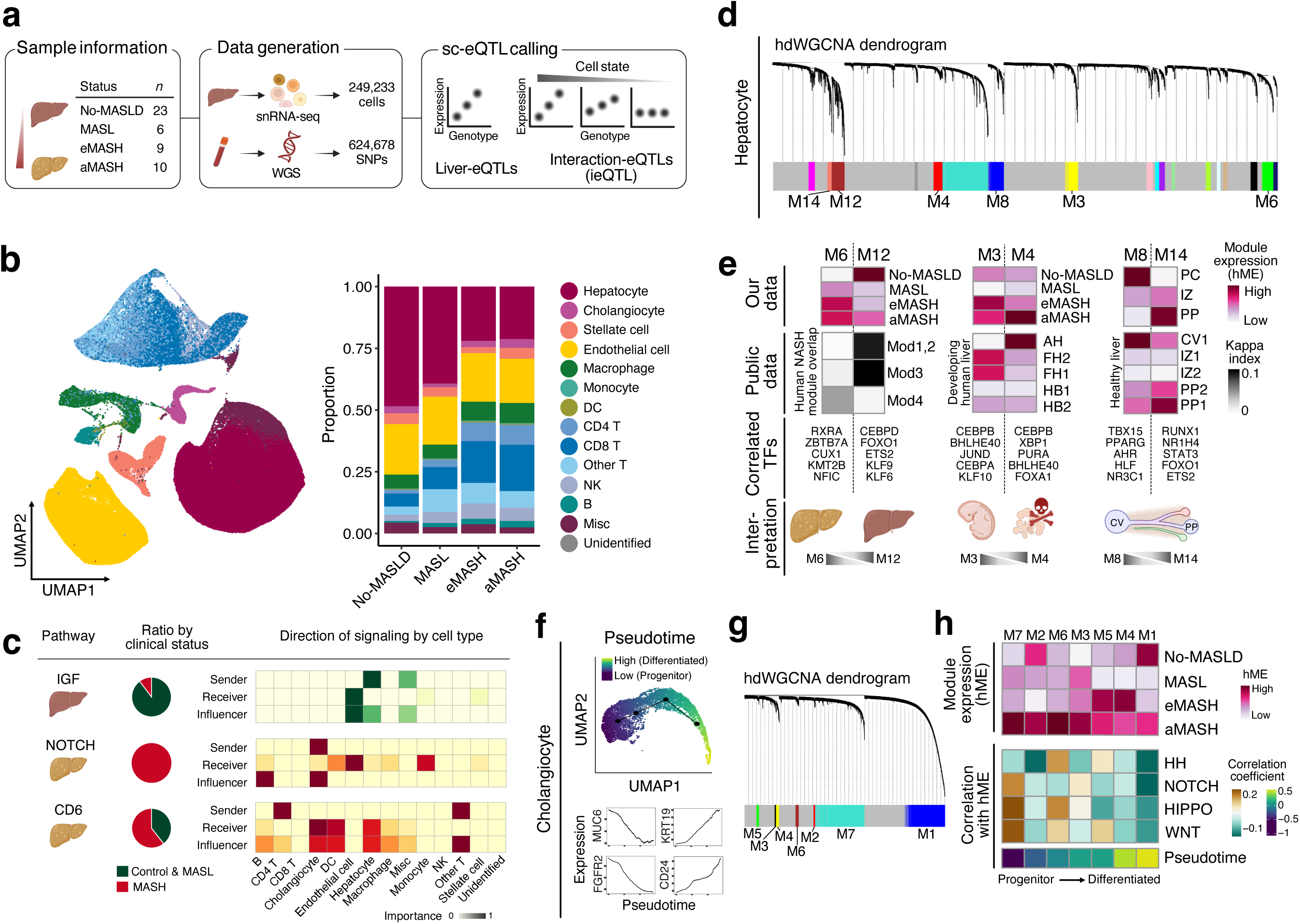
snRNA-seq reveals transcriptomic alterations within and across cell types in MASLD. **a**, Study scheme for identifying sc-eQTLs including liver-eQTLs and ieQTLs. **b**, Clustering of major liver cell types. UMAP plot showing major liver cell types (left) and their proportions by disease stage (right). **c**, Cell-cell interaction analysis. Pathways that are significantly down- (IGF) or up-regulated in MASH (NOTCH, CD6) are shown (left). Pie plot of relative information flow in non-MASH and MASH cells, computed by rankNet function in CellChat (middle). Heatmap displaying centrality scores for signaling roles (sender, receiver, and influencer), with green indicating scores from control cells and red from MASLD cells (right). **d**, Dendrogram from the hdWGCNA run performed on hepatocyte gene expression profiles. Each branch of the dendrogram represents a gene, and are grouped by colors to denote different gene modules. **e**, Characterization of hdWGCNA gene modules in hepatocytes. The top row shows heatmaps of harmonized module expression (hME) in our dataset. The second row shows heatmaps of hME of corresponding modules when projected to the public datasets. The third row displays top TFs whose activity correlated with hME. The bottom row illustrates the biological interpretation of each module. **f**, Pseudotime analysis in cholangiocytes. UMAP plot colored by pseudotime values (top). Expression of progenitor-associated genes (*FGFR2* and *MUC6*) and differentiated cell-associated genes (*KRT19* and *CD24*) along the pseudotime axis (bottom). **g**, Dendrogram of gene modules identified by hdWGCNA in cholangiocytes. **h**, Heatmap showing the module expression in each disease stages (hME; upper). Heatmap displaying the correlation coefficients of hME with developmental pathway genes expression (middle) and with pseudotime (lower).

Gene expression profiling of each cell type yielded specific marker gene expression patterns that aligned with a previous normal liver snRNA-seq study^43^ (Supplementary Methods and Supplementary Fig. 3). Notably, approximately half of the hepatocytes in the MASH livers were replaced with immune cells (hepatocyte proportion: 48.4% in no-MASLD vs. 21.2% in aMASH), reflecting increased inflammation and immune cell infiltration alongside hepatocellular death in the injured livers (Fig. 1b). Hepatocyte proportion was also inversely correlated with serum ALT, AST, HOMA-IR, NAFLD activity score, and NAFLD fibrosis score, further confirming that our experimental approach accurately reflects the clinicopathologic changes in patient’s liver tissue (Supplementary Fig. 4),

Cell type-specific gene expression varied across the disease phenotype groups, displaying enrichment of various MASLD-associated biological processes consistent with previous observations (*e.g.*, increased lipid and cholesterol metabolism and ubiquitin-mediated proteolysis; decreased drug catabolic processes for hepatocytes)^44,45^ (Supplementary Table 3).

Ligand-receptor-mediated intercellular communications are pivotal in maintaining tissue homeostasis and responding to injury. We observed several notable cell-cell interactions (Fig. 1c and Supplementary Fig. 5); for example, IGF signaling, an inhibitory signal against inflammation and fibrosis originating from the hepatocytes^46^ ^47^, acts on endothelial cells and is downregulated in the MASH liver (Fig. 1c). Meanwhile, Notch signaling, originating from cholangiocytes, acts on endothelial and immune cells and is elevated in the MASH liver, consistent with previous studies^48,49^ (Fig. 1c). Finally, CD6 signaling from CD4 T cells to hepatocytes, previously reported in autoimmune hepatitis^50^, is also activated in MASH (Fig. 1c). Thus, our snRNA-seq analysis revealed transcriptomic alterations within and across different cell types in MASH.

### Gene modules captured from each cell type illustrate cellular programs involved in liver cell stress, regeneration, and zonation

The expression levels of functional gene modules can be utilized as a proxy for specific cell states or cellular programs activated in individual cells. This approach enables more realistic simulation of *in vivo* cell status than the traditional binary categorization (“healthy” vs “diseased”). Through co-expression network analysis across the broad spectrum of MASLD cells^51^, we identified a total of 37 modules from four major cell types (hepatocytes, cholangiocytes, hepatic stellate cells, and endothelial cells; Supplementary Table 4).^43^ We focused our analysis on these cell types since snRNA- seq allows better expression profiling for these cells and they are functionally important in the liver.

We then investigated the biological implications of each module by comparing module expression profiles among MASLD disease stages, projecting them into public datasets (Supplementary Table 5 and Supplementary Fig. 6), and discovering transcription factors (TFs) whose activity correlates with module expression.

In hepatocytes, we identified six modules associated with cell stress, regeneration, or hepatic zonation patterns (Fig. 1d). Hepatocyte module 6 (Hep-M6) was upregulated in MASH cells in our cohort, and its network was well preserved in a recent human MASH snRNA-seq study that identified MASH-related gene modules (Mod1-4)^52^ (Fig. 1e, left). Hep-M6 overlapped with the MASH module (Mod4) representing reduced metabolic function and augmentation in cell adhesion and migration. Conversely, hepatocyte module 12 (Hep-M12) exhibited decreased expression in MASH and overlapped with modules from hepatocytes undergoing metabolic adaptation (Mod1-3; Fig. 1e, left).

Hep-M12 expression was strongly correlated with the activity of the TF FOXO1 (*R* = 0.63, *P* < 2.2 x 10^-308^) (Fig. 1e, left), which regulates glucose, triglyceride, and cholesterol homeostasis^53^. Similarly, Hep-M3 and Hep-M4 are related to regeneration and are preserved in the developing human liver (Fig. 1e, middle), while Hep-M8 and Hep-M14 are zonation-associated modules conserved in the normal human liver (Fig. 1e, right).

We likewise captured seven gene modules from cholangiocytes and correlated their expression with a pseudotime gradient. Since this gradient well captured the heterogeneity of cholangiocytes along the axis of progenitor to differentiated states^43,54^ (Fig. 1f), we also examined specific gene expression modules for implications in liver development (Fig. 1g, Supplementary Methods and Supplementary Table 6). Notably, cholangiocyte module 7 (Chol-M7) was highly expressed in cholangiocytes from advanced MASH patients but displayed an inverse correlation with the pseudotime gradient (*R* = −0.56, *P* = 2.2 x 10^-16^) and thus represents a progenitor cholangiocyte state. This module was positively correlated with Notch, Hippo, and Wnt signaling pathways (Fig. 1h). Other cholangiocyte modules showed varying degrees of correlations with the pseudotime gradient and developmental pathways (Fig. 1h), implying the presence of complex gene programs in cholangiocytes.

Additional analysis on hepatic stellate cells and endothelial cells also allowed us to select biologically interpretable gene modules in each cell type (Supplementary Tables 5 and 7). We considered the expression profiles of those modules as proxies for specific cell states in subsequent analyses, including interaction testing for ieQTL identification.

### Single-cell-eQTL mapping detects MASLD-associated gene regulatory variants

To understand variants that regulate hepatic gene expression, we mapped liver-eQTLs using the PME model at single-cell resolution. While many sc-eQTL studies have employed a linear model (LM) with pseudobulk aggregation, the PME model has the advantage of accounting for heterogeneity between single cells and hence simulating the complex physiology of the human liver. We followed a previously published method with modifications^37^ and identified a total of 3,553 liver-eGenes from four cell types (Fig. 2a). Most of the liver-eQTLs were cell-type-specific (58.3–72.8%; Fig. 2a), indicating distinct gene regulatory landscapes.

**Figure 2.**
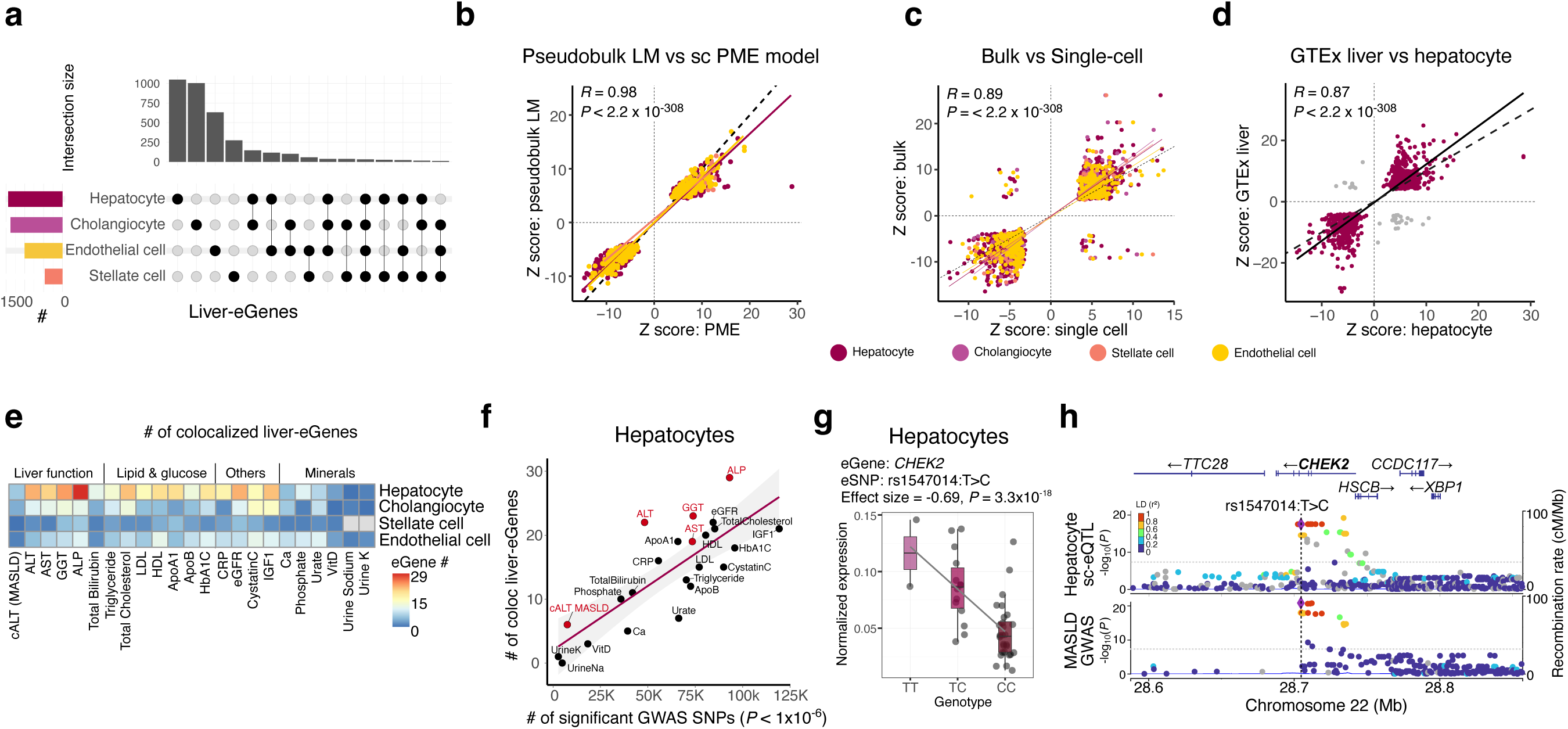
MASLD-associated gene regulatory variants detected by sc-eQTL mapping. **a**, Upset plot illustrating the count of significant liver-eGenes across four major cell types. **b**-**d,** Replication of liver-eQTLs using **b**, a pseudobulk LM , **c**, bulk-eQTL data from a different MASLD cohort, and **d**, GTEx liver data. Correlations were calculated and plotted using normalized effect sizes (Z-score = β/standard error) of significant gene-SNP pairs. **e**, Heatmap depicting the number of liver-eGenes that colocalized with various GWAS variants. **f**, Scatter plot displaying the relationship of the number of colocalizing hepatocyte liver-eQTLs (y-axis) with the number of significant GWAS SNPs from UK Biobank phenotypes (x- axis). The line of best fit is shown, with the 95% confidence interval in grey. Liver enzyme traits are highlighted in red. **g**, eQTL plot showing *CHEK2* expression in hepatocytes across genotypes of the MASLD-associated variant rs1547014:T>C. Expression of *CHEK2* was averaged per donor for visualization. **h**, Regional plot demonstrating the colocalization of hepatocyte liver-eQTL and MASLD GWAS variants.

To test the robustness of this method, we compared the standardized effect sizes of our significant liver-eQTLs with the effect sizes calculated from (i) the pseudobulk-LM, (ii) bulk eQTL from a different MASLD cohort, and (iii) GTEx liver eQTLs. As expected, significant liver-eQTLs showed highly strong correlations of effect sizes (*R* = 0.98, 0.89, and 0.87 for pseudobulk eQTLs, bulk eQTLs, and GTEx liver eQTLs, respectively; Fig. 2b-d and Supplementary Fig. 7).

To identify the clinical implications of the identified liver-eQTLs, we obtained GWAS summary statistics^55^ for traits relevant to MASLD and tested for the presence of variants having both GWAS and eQTL signals (*i.e.*, colocalized signals). Multiple liver-eQTLs colocalized with GWAS loci for liver function-related (*i.e.*, serum ALP, GGT or ALT) and metabolism-associated phenotypes (*i.e.*, total cholesterol or LDL levels) (Fig. 2e, Supplementary Fig. 8, and Supplementary Table 8). Most colocalizing liver-eQTLs originated from hepatocytes, and the number of colocalizing liver-eQTLs for each liver-associated phenotype exceeded the expected number (standardized residuals for ALT = 2.2, ALP = 1.8; Fig. 2f). An example of hepatocyte-originated liver-eQTL colocalizing with a recent MASLD GWAS locus^8^ is illustrated by the gene *CHEK2* and its eSNP rs1547014 (Fig. 2g). *CHEK2* is an important regulator of DNA damage response and also a suppressor of hepatocarcinogenesis^56^. This MASLD-associated variant is located in an intron, with the C allele being the risk allele (Fig. 2h)^8^; carriers of this allele exhibit decreased expression. This same variant is also an eQTL for liver and thyroid bulk tissues in GTEx (Supplementary Fig. 9).

### eQTLs interaction with cell states confers an additional layer of liver function regulation

As snRNA-seq allows us to document the heterogeneity of individual cells, we sought to incorporate cell condition information when assessing variant effects on gene expression regulation and disease association. This yielded a set of dynamic eQTLs that interacted with specific cell states (ieQTL; Fig. 3a). Cell states were defined as quantitative measurements of the expression levels of biologically interpretable modules (*i.e,* hdWGCNA modules) or as a binary value of disease status. Additionally, to exclude variants with no genotype effect, we restricted the interaction analyses to significant eQTLs identified from all donor cells (*i.e.*, the liver-eQTLs described in Fig. 2), MASLD cells, or control cells (Fig. 3a). Among a total of 10,507 eGenes identified from four cell types, 2,136 eGenes had significant ieQTLs (Fig. 3b and Supplementary Fig. 10).

**Figure 3.**
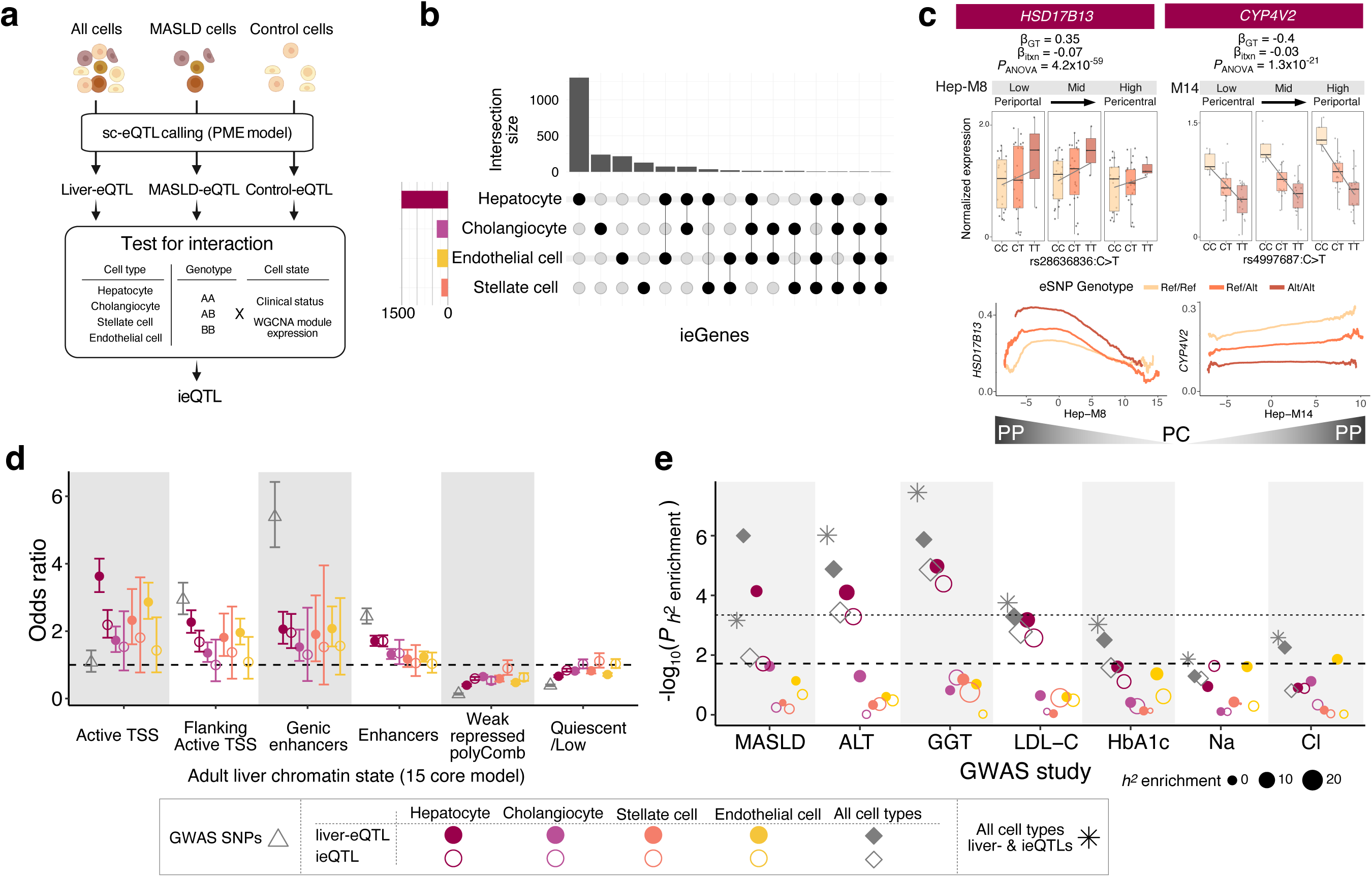
Interaction of eQTLs with cell states confers an additional layer of liver function regulation. **a**, Scheme for identifying ieQTLs. Broad set of sc-eQTLs were separately identified by the PME model among all cells (*i.e.* liver- eQTLs), MASLD cells, and control cells. For significant eQTLs, interactions of genotype with cell states were tested to identify ieQTLs. **b**, Upset plot of significant ieGenes from four major cell types. **c**, Examples of hepatocyte ieQTLs in *HSD17B13* and *CYP4V2* that exhibit significant interaction with the inferred zonal location of cells. Boxplots in the upper row show gene expression according to genotype and zonal location as inferred from module expression bins. Lower row shows kernel regression lines of expression by each genotype over module expression, which indicate distinct zonation patterns. PC, pericentral; PP, periportal region **d**, Dotplot displaying the odds ratio (OR) enrichments of sc-eQTLs in different chromatin states with 95% confidence intervals calculated using Fisher’s exact test. Dashed line: OR = 1. **e**, Heritability enrichments of sc-eQTLs in various phenotypes from a Japan BioBank GWAS, calculated by S-LDSC. Dot size denotes the enrichment of heritability relative to what would be expected by chance. Dashed and dotted lines respectively indicate significance cutoffs for *P* = 0.05 or *P*_adj_ = 0.05 adjusted by the Benjamini and Hochberg method.

These interacting-eGene (ieGenes) were not particularly enriched in any specific module (mean phi correlation coefficient = 0.042; Supplementary Fig. 11). The module most abundantly represented among eGene interactions was Hep-M8 (pericentral- hepatocyte-specific module; 2,859 eGenes), followed by Hep-M14 (periportal- hepatocyte-specific module; 2,807 eGenes; Supplementary Fig. 12). One notable example is an ieQTL in *HSD17B13*, whose inhibitory influence exerts a protective effect against MASLD and liver fibrosis^14,57^. The effect size of the eQTL was elevated in periportal zones, with the highest *HSD17B13* expression observed in individuals carrying the rs28636836:TT genotype (Fig. 3c). This eQTL was not found significant in the bulk liver, demonstrating the higher sensitivity of the ieQTL approach toward context- and genotype-specific signals (Supplementary Fig. 9). Another eQTL specific to the periportal region is in *CYP4V2*, an emerging druggable target for MASLD due to its association with fatty acid metabolism^58^. Expression of this gene was increased in the periportal region of the CC genotype carriers (Fig. 3c). These findings highlight the utility of ieQTLs by uncovering genotype- and zone-specific regulation of previously identified disease-associated genes in the liver.

To systematically evaluate the genetic implications of ieQTLs along with liver-eQTLs, we annotated eSNPs with epigenetic states and assessed their enrichment. Our eSNPs were enriched with promoter states (hepatocyte eSNPs OR = 3.63, *P* = 2.01 x 10^−56^; Fig. 3d) while GWAS SNPs were enriched with enhancer states (OR = 5.4, *P* = 1.98 x 10^−51^; Fig. 3d), which is in line with a previous observation^21^. In general, ieQTLs exhibited weaker enrichments in each chromatin state compared to liver-eQTLs (Fig. 3d), likely due to the context-specific nature of ieQTL effects. Furthermore, stratified LD score regression analysis showed that liver-eQTLs and ieQTLs are significantly enriched with MASLD heritability (liver-eQTLs: OR = 5.39, *P* = 9.99 x 10^−7^; ieQTLs: OR = 10.27, *P* = 1.21 x 10^−2^; Fig. 3e) and liver enzyme levels (for ALT levels, liver-eQTLs: OR = 10.42, *P* = 1.32 x 10^−5^; ieQTLs: OR = 13.9, *P* = 3.74 x 10^−4^; Fig. 3e). While enrichment *P*-values were stronger for liver-eQTLs (Fig. 3e), enrichment estimates were higher in ieQTLs (Supplementary Table 9) for ALT, AST, GGT, and other traits.

Ultimately, it can be concluded that liver-eQTLs and ieQTLs are in epigenetically active regions and contribute to phenotypic variance in liver function.

### An integrative analysis of ieQTLs, their upstream regulators, and phenotypes provides mechanistic insights into regulatory variants

The most plausible mechanism of eQTL action is through conferring differential affinity to upstream TFs that can regulate eGene expression (Supplementary Fig. 13)^19,59^. Thus, we hypothesized the existence of a “master” upstream regulator of cell-state-specific ieQTLs, which would be functionally implicated in disease progression (Fig. 4a)^60^. TF activity was inferred from single-cell data using pySCENIC^61,62^, and the creation or disruption of TF binding motifs by interacting-eSNP (ieSNP) presence was assessed, as such changes can provide a mechanistic explanation of cell-state-specific ieQTLs (Fig. 4b). This analysis yielded 601 TF-cell state-ieSNP-ieGene “quartets” from the four cell types. Among the 601 quartets, a majority was found in hepatocytes (506/601 = 84.2%). In the hepatocyte-originated quartets, Hep-M14 (represents periportal module) was found in 238 quartets and Hep-M12 (represents healthy hepatocyte status) was associated with 167 quartets. TFs and ieGenes that are parts of the Hep-M14 quartets also displayed a periportal-biased expression pattern (Supplementary Fig. 14a) and ieGenes that are components of the Hep-M12 quartets were enriched in various metabolic pathways active in the liver (Supplementary Fig. 14b). Twenty-three of the 601 quartets were significantly associated with MASLD or liver phenotypes, as their ieSNPs were also found to be significant in the GWAS using the same traits^8,55,63^ (Fig. 4b, c and Supplementary Table 10). Among the 23 quartets, KLF6 and EGR1 were the most frequently appearing TFs (Supplementary Table 10). Notably, the strongest cell state-TF pair correlation was between Hep-M12 and FOXO1 (Supplementary Table 10). Among the potential transcriptional targets of FOXO1, *EFHD1* expression and rs13395911 genotype showed a significant interaction with Hep- M12, which was also corroborated by the presence of bulk-eQTL from GTEx liver tissue (Fig. 4d-1). The eQTL effect diminished in cells with low Hep-M12 expression (Fig. 4d- 1), characteristic of diseased hepatocytes (Fig. 1e and Fig. 4d-2). Hep-M12 expression also correlated with FOXO1 activity (Pearson *R* = 0.63; Fig. 4d-3), which was similarly decreased in cells from MASLD donors (Fig. 4d-4,5). A new FOXO1 binding motif was created by the alternative allele of rs13395911 (*P* = 7.01 x 10^−5^ from the FIMO software; Fig. 4d-6), and a statistically significant correlation of *EFHD1* expression with *FOXO1* expression at a single hepatocyte level was shown in donors with the TT genotype (linear regression *P* = 0.01), but not in those with either the AT genotype (linear regression *P* = 0.94) or AA (linear regression *P* = 0.51) genotype, implying an allele-specific regulatory effect of *FOXO1* expression on *EFHD1* (Supplementary Fig. 15). The variant colocalizes with the signal from a serum ALT GWAS^63^ (Fig. 4d-7), and is in strong LD with a MASLD GWAS hit rs7604422^8^ (*R^2^* > 0.9). These results suggest that the regulatory effect of a MASLD risk locus on *EFHD1* expression is mediated by FOXO1, linking it to genotype-specific disease susceptibility (Fig. 4e). More specifically, in rs13395911:AA donors, homozygous for the protective allele (54.8% in East Asian populations, 16.3% in European, and 13.4% in African), FOXO1 binding would be weak and *EFHD1* expression remains low regardless of the cell state (Hep-M12). Meanwhile, in TT donors harboring the risk allele (6.8% in East Asian populations, 35.5% in European, and 40.2% in African), FOXO1 strongly binds to the eSNP region and activates *EFHD1* expression. Indeed, decreased FOXO1 in maladapted hepatocytes leads to the loss of *EFHD1* expression (*EFHD1* mean expression fold change between Hep-M12 highest quartile vs lowest quartile in MASLD cells = 1.56; Supplementary Fig. 16). Although a previous study reported elevated FOXO1 activity in human MASH^64^, our snRNA-seq and bulk RNA-seq results^29^ (Fig. 4d-5,6, Supplementary Fig. 17) consistently point to decreased FOXO1 activity in MASH livers, in accordance with the results from mouse studies^65–67^. These observations generated a hypothesis that during hepatocyte maladaptation, reduced FOXO1 expression induces the dynamic downregulation of *EFHD1* in TT donors, which may foster MASLD.

**Figure 4.**
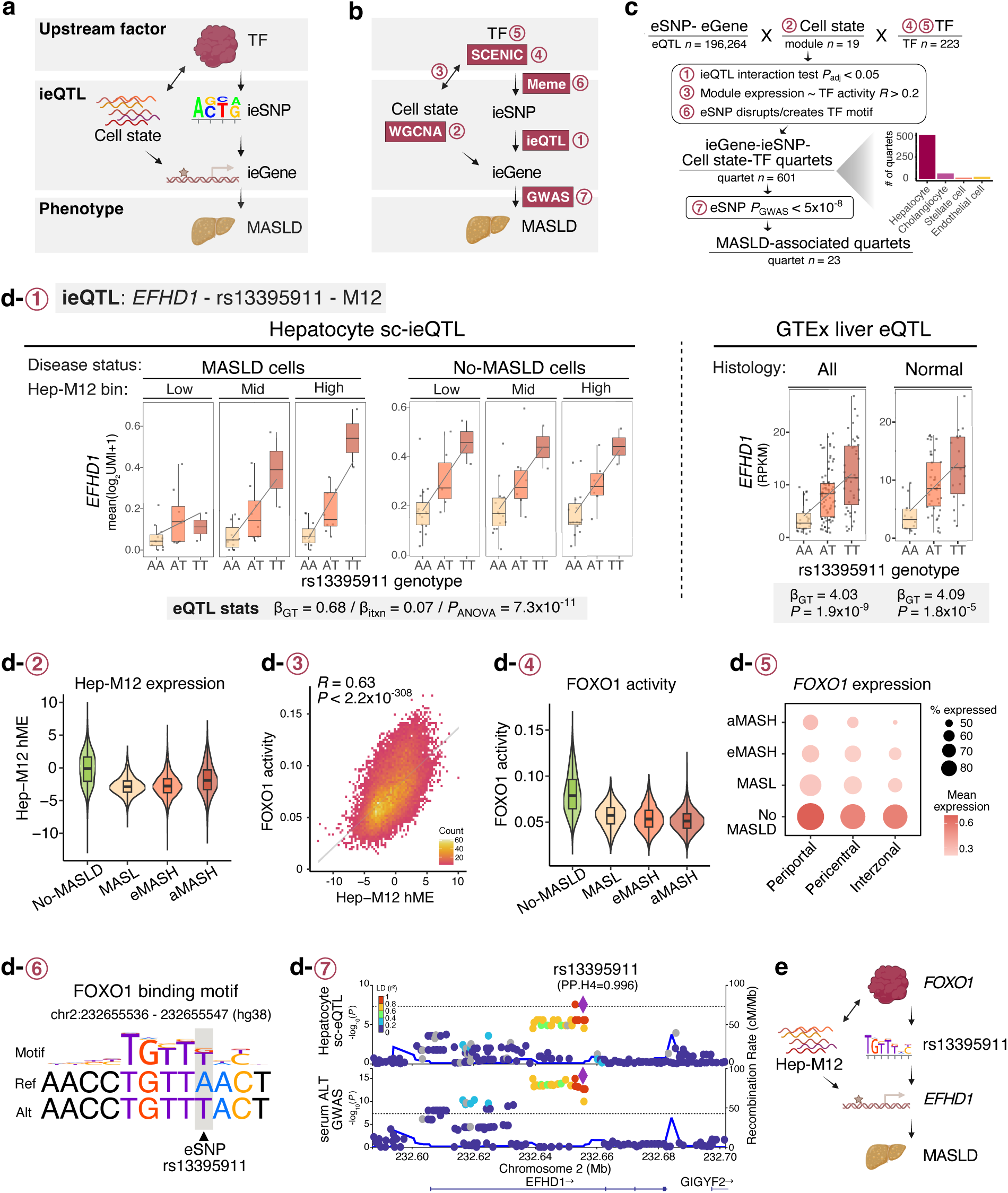
Integrative analysis of ieQTLs, upstream regulators, and phenotypes identifies genotype-specific regulatory variants contributing to MASLD. **a**, Schematic illustration of the relationship between upstream TFs, ieQTL components (cell states, ieSNPs, and ieGenes) and the phenotype (MASLD). **b**, Pipeline for prioritizing MASLD-associated regulatory variants and their upstream factors. Purple boxes highlight the method used in each process. **c**, Analysis pipeline scheme with the count of prioritized quartets (TFs, cell states, ieSNPs, and ieGenes) at each step. **d**, Example of a MASLD-associated ieQTL in *EFHD1* and its upstream regulator *FOXO1*. Subplots (1-7) detail each step in the analysis pipeline from Fig. 4b: **d-1**, eQTL plot for an ieQTL in *EFHD1* plotted against genotype of rs13395911 (chr2:232655544:A>T) in cells derived from MASLD donors (left), control donors (middle), and GTEx bulk liver (right). Regression coefficients (ß_GT_ and ß_itxn_ indicate PME coefficients for the genotype and interaction term, respectively) and *P*-values (*P*_ANOVA_ was obtained from the interaction test) are displayed below. **d-2**, Expression of Hep-M12 across disease stages, showing reduction in MASLD. **d-3**, 2D histogram showing correlation of Hep- M12 expression with FOXO1 activity in hepatocytes. **d-4**, Violin plot of FOXO1 activity across disease stages. **d-5**, Dotplot of *FOXO1* expression across disease stages and hepatocyte clusters. **d-6**, Alignment of FOXO1 motif (sequence logo) with eSNP sequences. **d-7**, Regional plot of the sc-eQTL in *EFHD1* and serum ALT GWAS from Japan BioBank indicating the probability of colocalization (PP.H4 on top). **e**, Schematic representation of the ieQTL in *EFHD1*, rs13395911, FOXO1 and its link to MASLD.

**Figure 5.**
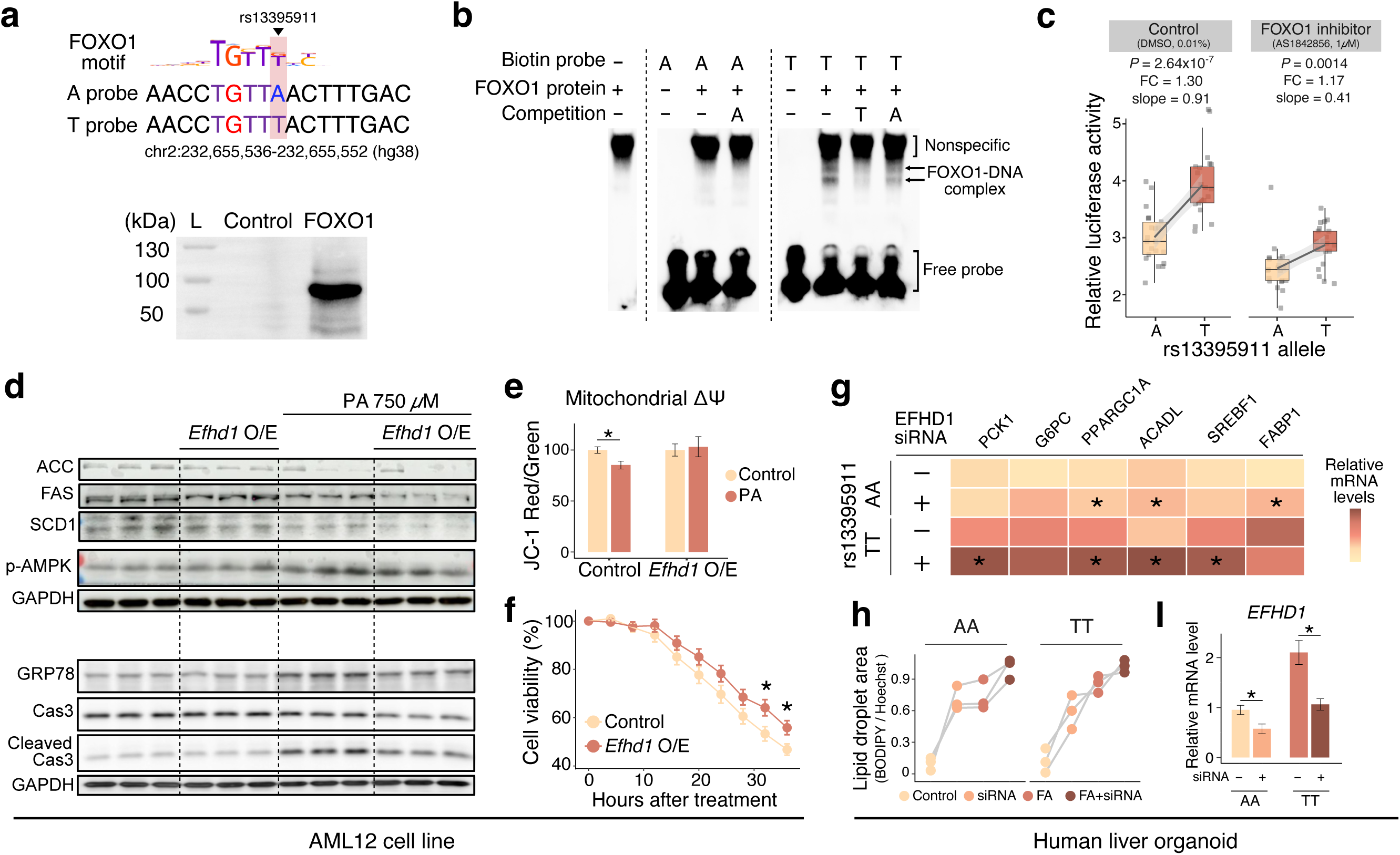
Experimental validation of the FOXO1-rs13395911-EFHD1-MASLD axis. **a**, DNA probes used for EMSA (top). Western blot for the *in vitro*-translated FOXO1 protein (bottom). **b**, EMSA result indicating T allele-specific FOXO1 binding. Competition assays were conducted with a 10x molar excess of cold probes. **c**, Luciferase reporter assay showing the differential transcriptional activity between A and T alleles, which was attenuated by FOXO1 inhibitor treatment. *P*-values, fold changes (FC) between alleles, and slopes from linear regression are annotated above the plot. **d**, Western blot of proteins involved in lipid synthesis (upper), ER stress, and apoptosis (lower) in AML12 cells **e**, Mitochondrial membrane potential (Δψ) measured using JC-1 dye in AML12 cells. Mean values with error bars representing standard error are shown. **f**, Cell viability in AML12 cells following tunicamycin treatment. Mean values with error bars representing standard errors are shown. **g**, Relative mRNA levels of genes associated with glucose or lipid metabolism. *, *P-*value < 0.05 (t-test). **h**, Lipid droplet accumulation of hepatic organoids measured as ratio of area of green fluorescence (BODIPY) by blue fluorescence (Hoest33342). Data points obtained from the same iPSC-derived organoids are connected with a gray line. **i**, Relative mRNA levels of *EFHD1*. Mean values with error bars representing standard error are shown. *, *P-*value < 0.05 (t-test). Abbreviations: O/E, overexpression; FC, fold change; FA, free fatty acid treatment

### The MASLD risk locus rs13395911 exerts a regulatory effect via FOXO1 on EFHD1 expression in pathologic cell states

To validate the functional implication of the FOXO1-maldaptive state-rs13395911-*EFHD1* axis, we first confirmed the differential binding activity of FOXO1 on the DNA element flanking the ieSNP using the electrophoretic mobility shift assay (EMSA) (Fig. 5a, b). Then, we confirmed the transcription regulatory effect of rs13395911 through FOXO1 in HepG2 cells using a reporter assay (Fig. 5c, two-sided t-test *P* = 2.64 x 10^−7^). This effect was diminished when cells were treated with a FOXO1 inhibitor (Fig. 5c, slope changed from 0.91 to 0.41 by inhibitor addition). Therefore, T-allele-specific FOXO1 binding at rs13395911 regulates the expression of *EFHD1*.

Next, we sought to understand the molecular function of EFHD1 in hepatocytes. A knockdown assay was not feasible as the baseline expression of *EFHD1* was indiscernible in mouse hepatocyte cell line (AML12). Instead, we overexpressed *Efhd1* in AML12 and observed decreased expression of proteins related to lipid synthesis (FAS and SCD1), endoplasmic reticulum (ER) stress (GRP78), and apoptosis (cleaved Cas3) (Fig. 5d). *Efhd1* overexpression also de-repressed the mitochondrial membrane depolarization caused by palmitic acid (PA), a sign of early apoptosis (Fig. 5e), and restored the viability of cells treated with the ER stress inducer, tunicamycin (Fig. 5f). These results suggest that EFHD1 has a protective role against lipotoxic processes in hepatocytes.

Then we sought to demonstrate the genotype-dependent protective effect of EFHD1 *in vivo*, which would not be feasible in the mice as rs13395911 is located in the deep- intronic region of *EFHD1* and the FOXO1-binding element is not conserved in rodents (Supplementary Fig. 18). Thus, human hepatic organoids harboring different genotypes in rs13395911 (three with rs13395911:AA and three with rs13395911:TT; Supplementary Table 11) were established and analyzed.^68,69^ Knocking-down *EFHD1* in the organoids increased expression of gluconeogenesis genes (*PCK1, G6PC,* and *PPARGC1A*) and lipid metabolism genes (*FABP1, SREBF1,* and *ACADL*) and enhanced accumulation of intracellular lipid droplets, mimicking the MASLD phenotype (Fig 5g, h, and Supplementary Fig. 19). Notably, we observed genotype-specific effects of *EFHD1* inhibition, such as TT-genotype specific up-regulation of *PCK1* and *SREBF1*^70^, and AA-genotype specific up-regulation of *FABP1*, a cytoprotective gene against lipotoxicity^71,72^ (Fig. 5g). These findings suggest that loss of *EFHD1* expression contributes to exacerbation of MASLD, particularly in rs13395911:TT donors. Moreover, *EFHD1* siRNA-treated rs13395911:TT-genotype organoids, which displayed characteristics of metabolic dysfunction, had *EFHD1* levels comparable to those of control siRNA-treated AA-genotype organoids (*EFHD1* siRNA-TT/control siRNA-AA fold difference = 1.12, *P* = 0.46; Fig. 5i). This observation suggests that the relative change in the level of *EFHD1* plays a critical role in the pathogenic process.

Taken together, decreased FOXO1 leads to the dynamic loss of *EFHD1* expression in TT donors, resulting in aggravated ER stress, mitochondrial dysfunction, apoptosis, and metabolic dysregulation, all of which contribute to MASH progression.

## Discussion

MASLD is a heterogeneous and heritable disease with many clinically relevant genes and variants. Simultaneous profiling of single-cell transcriptomics and inherited genetic variation opens opportunities to elucidate the complex interactions of these genetic factors. Our comprehensive analysis of single nuclei from human liver tissues unraveled the intricate interplay among genetic variation, gene expression, and cell state at single-cell resolution. We delineated genetic effects on gene expression in four major liver cell types, uncovering more than 3,500 *cis*-acting liver-eQTLs. Moreover, we suggest a sc-eQTL-based framework that allows for the discovery of functional units comprised of interacting TFs, cellular states, ieSNPs, and ieGenes (*i.e.*, quartets). We demonstrated that these loci are associated with MASLD and explain a substantial proportion of phenotypic variability.

Zonation in the liver disparately controls the function and state of individual cells, variably affecting MASLD patients^73^. In our analysis, genetic regulation also exhibited a specific zonal pattern, with zonation predicting the most significant ieQTLs among the cell states tested. One example is the transcriptional regulation of *HSD17B13* by rs28636836 in the periportal region (Fig. 3c). The observation that *HSD17B13* is highly expressed by periportal hepatocytes in rs28636836:TT donors has implications for patient stratification in ongoing clinical trials using RNA-based therapeutics and for the discovery of novel zonation- and genotype-specific treatment targets.

Additionally, we propose that rs13395911 increases the risk of MASLD through allele-specific FOXO1 binding and modification of *EFHD1* expression (Fig. 4, 5). EMSA and reporter assay validated the FOXO1-rs13395911-*EFHD1* axis, implicating it as a potential genotype-dependent personalized therapeutic target. In line with our discovery, a recent study identified a rs13395911 allele-specific opening of chromatin at the *EFHD1* region through differential interaction with FOXA2^74,75^. We did not prioritize FOXA2 due to its negative correlation with cell state (Hep-M12; *R* = −0.23). However, given that FOXO1 and FOXA2 bind to DNA interdependently to remodel chromatin^76^, and considering that rs13395911 resides in an active enhancer element with various TF binding peaks^74^, it would be valuable to study the pleiotropic effects of this locus that can be mediated by the two TFs.

Based on functional studies in cell lines and hepatic organoids, we suggest *EFHD1* as a genotype-dependent protective gene in MASLD hepatocytes. In hepatocytes, we found EFHD1 to be intricately involved in lipid metabolism, ER stress, mitochondrial dysfunction, and apoptosis. Unlike EFHD2, which is expressed in hepatic myeloid cells and exerts a pro-inflammatory effect in MASH^77^, EFHD1 is known to play roles in reducing pathogenic mitochondrial calcium influx^78^ and maintaining mitochondrial membrane potential^79^, which enhances mitochondrial function in toxic environments.

Furthermore, recent studies revealed the influence of EFHD1 on calcium homeostasis varies among different cell types based on its baseline expression level^78^. Variable baseline *EFHD1* expression linked to rs13395911 genotypes might underlie the genotype-specific effects of EFHD1 loss during MASLD progression.

Due to the polygenic nature of MASLD and the substantial heritability enrichment found within sc-eQTLs, it is anticipated that more loci will merit evaluation for biological implications.

Our study utilized a large single-center East Asian cohort representing a wide spectrum of MASLD phenotypes, from isolated steatosis without fibrosis to advanced MASH with fibrosis stage ≥3. Furthermore, matched genotype data and our application of the PME model enabled joint analysis of genome and transcriptome in the disease context, revealing disease-associated regulatory variants and their mechanistic aspects.

The limitations of the current study include its small sample size, the limited number of genetic variants tested, and limited characterization of cell states. Employing sample multiplexing techniques and increasing the throughput of scRNA-seq would ultimately lead to a well-powered liver sc-eQTL study. Also, we restricted our eQTL analysis to the variants with the highest allele frequency within an LD block due to the computational time and multiple testing burden. This potentially underestimates the full range of eQTLs and hampers causal variant discovery. Future studies would benefit from faster and scalable computational resources and better statistical methods to control false positive rates in testing multiple hypotheses.

To summarize, this study integrated genetic variation with snRNA-seq to uncover genetic contributors to inter-individual differences in MASLD. Our findings illustrate how distinct genetic variations impact the expression of MASLD-related genes in a context-dependent manner and propose a novel mechanistic aspect underlying MASLD pathogenesis. Further elucidating the genetic foundations of liver cell states holds significant implications for patient stratification in clinical trials and the discovery of personalized therapeutic options.

## Acknowledgements

We thank the participants for sharing their valuable tissues. We also thank Kazuyoshi Ishigaki for assisting PME model adaptation, Hanbyul Kim and Chul-Hwan Lee for providing guidance in performing EMSA, and Juyoung Lee in mathematical modeling process. This work was in part supported by grants through the National Research Foundation (NRF) funded by the Ministry of Science & ICT (2021R1A2C2005820 and 2021-M3A9E4021818 to W.K. and 2023-00207857 and 2021-R1A2C3014067 to M.C.) and by a grant of the MD-PhD/Medical Scientist Training Program through the Korea Health Industry Development Institute (KHIDI), funded by the Ministry of Health & Welfare (to S.H.).

## Data availability

DNA sequencing and single nucleus transcriptome data of this study were deposited in K-BDS under ID KAD2462523. Processed Seurat object and codes for the analysis will be available upon manuscript acceptance.

## Author information

### Contributions

S.H., M.C., W.K. conceived and designed the experiments and analysis. W.K., K.-S.S. performed frozen liver sample processing and clinical data collection. S.H. performed snRNA-seq data analysis, genome data processing, eQTL calling and downstream analyses. Y.J.L. performed experiments in AML12 cell lines under K.W.K.’s supervision.

S.J.M. generated and performed experiments on hepatic organoids under M.J.S.’s supervision. T.Y. analyzed bulk liver RNA-seq and genotype data. S.H., M.C., W.K., S.J.M., Y.J.L., K.W.K., M.J.S. wrote the manuscript. All authors read and approved the manuscript.

### Corresponding authors

These authors jointly supervised this work: Murim Choi, Won Kim

## Ethics declarations

The authors declare no conflicts of interest that pertain to this work.

## Online Methods

### Cohorts

This study was approved by the institutional review board of Seoul Metropolitan Government Boramae Medical Center (IRB No. 30-2021-23 and 20-2021-92). Frozen liver tissues were obtained from 31 patients with biopsy-proven MASLD and three healthy controls selected from a prospective Boramae MASLD cohort registry.

Additional 20 normal liver tissue samples were acquired from the living donor liver transplantation bank at Seoul National University Hospital (IRB No. C-1810-096-980). Inclusion and exclusion criteria are listed in the Supplementary Methods and baseline clinical characteristics of donors are provided in Supplementary Table 1. Study participants are all Korean, biopsied at age 17-78, and visited Seoul Metropolitan Government Boramae Medical Center or Seoul National University Hospital. All participants were informed of the study and provided written and signed consent.

Histological diagnosis, MASLD activity scoring and fibrosis staging was performed by a single pathologist (Supplementary Methods). Patients were categorized into no-MASLD, MASL and MASH. MASH was further classified by fibrosis severity into early MASH (eMASH; F0-2) and advanced MASH (aMASH; F3-4).

### Single nucleus RNA-sequencing data generation

Frozen liver tissues were processed to generate 10x Genomics-based snRNA-seq data in GENINUS (Seoul, Republic of Korea) using the previously reported protocol^43,80^. Tissues were homogenized, and isolated nuclei was counted using flow cytometry (BD: FACSMelodyTM Cell sorter). 10x Genomics chromium instrument and cDNA synthesis kit v3 were used to generate barcoded cDNA libraries. cDNA library quality was determined using an Agilent Bioanalyzer. Two paired-end sequencing with a read length of 100 bp was performed with Illumina Novaseq6000.

### snRNA-seq data analyses

Read processing, expression counting, batch correction, cell annotation, and differential gene expression analyses were conducted following the standard protocol (see Supplementary Methods). Briefly, reads were aligned, filtered, and counted by Cellranger^81^ and ambient RNA was removed by Cellbender^82^. Output files were converted into the Seurat object implement in Seurat 4^83^ and further analyzed in R (version 4.1.0). Liger^84^ was used to correct for batches and Seurat was used to cluster cells. Annotations were performed manually (for non-immune cells) or using Azimuth^83^ (for immune cells). During the processes, two samples with low gene counts, one PCA outlier, and three samples with low genome-transcriptome concordance tested by VerifyBamID^85^ were excluded. Differential expression analysis was performed with Seurat FindMarkers. Pseudotime analysis was performed with Slingshot^86^. Intercellular interaction was analyzed by CellChat^87^. Transcription factor activity was quantified with pySCENIC^61,62^. Gene ontology enrichment was tested with enrichR^88^. Finally, each cell type was subjected to co-expression network analysis using hdWGCNA package^51,89^. Gene modules were projected to public scRNA-seq datasets to seek for preservation and biological interpretation.

### Genome data generation

DNA samples were extracted from the whole blood or liver tissues and were subjected to 1x low-coverage WGS (Gencove, New York, NY). Procedures for variants calling, filtering, and LD clumping are described in Supplementary Methods.

### liver-eQTL and ieQTL calling

A stringent quality control on samples and genes for liver-eQTL calling was applied on the 48 samples. Among these, four samples with relatively low cell counts for major cell types were excluded for eQTL calling. Autosomal genes expressed in more than 50% of donors and 5% of all cells were included. Poisson mixed effects (PME) model was modified from a previous study^37^ (see Supplementary Methods). Covariates included disease status, scaled biopsy age, sex, library size (nCount_RNA), five gene expression PCs. *P*-values from tested *cis*-SNPs were adjusted with Benjamini-Hochberg process in hierarchical manner. To examine whether the relationship between gene expression and genotype changes depend on cell state, we added an interaction term between genotype and cell state. Cell states were defined as biologically interpretable module expression (continuous) or disease status of the donor (binary). Likelihood ratio test (ANOVA function in R) comparing the model with and without interaction term was used to calculate *P*-value. Specifically, χ^2^-statistic was calculated as −2 ×log (𝑙𝑖𝑘𝑒𝑙𝑖ℎ𝑜𝑜𝑑 𝑟𝑎𝑡𝑖𝑜), which was compared against χ^2^ distribution with one degree of freedom. Such test was performed to a broad set of significant eQTLs called from all cells (*i.e.*, liver-eQTLs that were described in Fig. 2), MASLD donor cells, and control donor cells.

### Colocalization analysis

The likelihood of eQTLs and GWAS sharing a causal variant was assessed using the coloc R package^90^ (coloc.abf function). We downloaded GWAS summary statistics of 23 and 21 serum or urine biomarkers from UK Biobank^55^ and BioBank Japan^63^, respectively. For each liver-eGene, we included *cis*-SNPs present in and GWAS summary statistics. Also, liver-eGenes with at least one *cis*-SNP with GWAS *P* < 1x10^-6^ was used for analysis. Posterior probability of H4 > 0.8 was assigned with significant colocalization. To account for the effect of LD clumping, we recalculated the effect size, *P*-values, and the colocalization probabilities of liver-eQTLs with all *cis*-SNPs without clumping (Supplementary Fig. 8). Visualization of colocalizing signals were plotted in the LocusZoom^91^ plot format. Linear regression of colocalized liver-eGene counts on significant GWAS SNP counts were used to calculate expected number of colocalizations and standardized residuals.

### Stratified LD score regression (S-LDSC)

LD score estimation and heritability partitioning were performed with S-LDSC^92^. East Asian GWAS summary statistics from BioBank Japan^63^ was used. The Supplementary Methods provide descriptions of functions and options employed.

### Integrative analysis of ieQTLs

Significant ieQTLs were used for integrated analysis. Regulon AUC scores calculated by pySCENIC^61^ were used as an estimate of transcription factor (TF) activity. To calculate TF activity and cell state correlation, Pearson’s correlation between AUC scores and harmonized module expression values was calculated. For TF binding motif analysis, nucleotide sequences flanking 15 bp on each side of eSNP were submitted to the Meme suite FIMO software^93^. Motif occurrences was determined by the FIMO *P* < 10^-4^. If TF motif occurs in the sequence with the reference allele, but does not in alternative allele, or vice versa, it was designated as an eSNP that creates or disrupts TF motif. "Quartets” composed of TF, cell state, ieSNP, and ieGene that satisfy following criteria were selected: (i) significant ieQTL, (ii) Pearson’s correlation coefficient between TF activity and module expression > 0.2, and (iii) ieSNP creates or disrupts TF motif. Then, we annotated five MASLD-associated GWAS summary statistics (serum ALT and AST GWAS from UK Biobank^55^, serum ALT and AST GWAS from Biobank Japan^63^, and MASLD GWAS^8^) to the ieSNPs of the quartets. Quartets in which the eSNPs showed significant association with any of the five traits were designated as MASLD associated quartet.

### Functional validation of FOXO1-rs13395911-EFHD1 axis

Procedures for performing electrophoretic mobility shift assay (EMSA), luciferase reporter assay, western blot, *Efhd1* overexpression, membrane potential measurement, cell viability monitoring, siRNA mediated knockdown in liver organoids are described in the Supplementary Methods.

