## Supplemental methods and figures for "Identification of genes conferring individual-level variation responsible for metabolic dysfunction-associated steatohepatitis using single-cell eQTL analysis"

**Supplementary Materials and Methods**

**Supplementary Figures**

Supplementary Figure 1. eQTL power analysis.

Supplementary Figure 2. snRNA-seq data quality control metrics

Supplementary Figure 3. Correlation with a public snRNA-seq study

Supplementary Figure 4. Correlation between cell type proportion and clinical features of patients.

Supplementary Figure 5. Differences in the overall cell-cell interaction patterns in No-MASH versus MASH cells.

Supplementary Figure 6. Reproducibility of hdWGCNA module networks in public data

Supplementary Figure 7. Replication of liver-eQTLs with other model or cohorts.

Supplementary Figure 8. Colocalization analysis without LD clumping on cis-SNPs

Supplementary Figure 9. Sc-eQTLs in GTEx dataset

Supplementary Figure 10. Counts of liver-eQTLs, MASLD-eQTLs, control-eQTLs and ieQTLs

Supplementary Figure 11. Module membership of ieGenes

Supplementary Figure 12. Upset plot of cell states that had significant ieQTLs

Supplementary Figure 13. eSNPs are enriched in known TF binding sites

Supplementary Figure 14. Analysis of quartets associated with Hep-M14 or M12 module-related cell states.

Supplementary Figure 15. Correlation between FOXO1 and EFHD1 expression

Supplementary Figure 16. EFHD1 expression fold change in different Hep-M12 quartiles in AA/TT donors.

Supplementary Figure 17. Loss of FOXO1 expression during MASLD progression

Supplementary Figure 18. Multiple alignments of sequences near rs13395911 in human, primates and rodents

Supplementary Figure 19. Lipid staining and RT-qPCR results from human hepatic organoids

Supplementary Figure 20. Comparison of various batch correction methods

Supplementary Figure 21. Genotyping accuracy in repeat regions

Supplementary Figure 22. Distribution and size of LD clumps

Supplementary Figure 23. Overexpression of Efhd1 in AML12 cells validated with RT-qPCR

**Supplementary Tables**

Supplementary Table 1. Baseline characteristics of the MASLD cohort according to disease group

Supplementary Table 2. Cell counts per cell type and donor

Supplementary Table 3. Gene ontology terms enriched among DEGs between no-MASH(ctrl & MASL) and MASH cells

Supplementary Table 4. hdWGCNA module genes for hepatocytes (4-1), Cholangiocytes (4-2), Stellate cell (4-3), and endothelial cells (4-4).

Supplementary Table 5. List of public single-cell data used for hdWGCNA module projection

Supplementary Table 6. Cholangiocytes related gene list

Supplementary Table 7. Modules representing specific cell state

Supplementary Table 8. List of colocalizing liver-eGenes

Supplementary Table 9. Heritability enrichment P value and estimates of eQTLs and ieQTLs.

Supplementary Table 10. List of quartets and their statistics.

Supplementary Table 11. iPSC lines used for hepatic organoid generation

Supplementary Table 12. Celltype marker genes

Supplementary Table 13. Oligonucleotide probes used for EMSA

Supplementary Table 14. Vectors used for luciferase reporter assay

Supplementary Table 15. Medium composition for maintenance and differentiation of hepatic organoids

Supplementary Table 16. List of RT-qPCR primers used in this study

### Supplementary Methods

*Cohort inclusion and exclusion criteria*

We constructed a prospective cohort from the ongoing Boramae MASLD registry (NCT02206841). Subjects with radiologic evidence of hepatic steatosis were eligible for study inclusion from January 2013. The eligibility criteria for this study were as follows: (i) ⩾18 years old, (ii) bright echogenic liver on ultrasound scanning (increased liver/kidney echogenicity and posterior attenuation), and (iii) unexplained high alanine aminotransferase (ALT) levels above the reference range within the past 6 months. The following exclusion criteria were used: (i) hepatitis B or C virus infection, (ii) autoimmune hepatitis, (iii) drug-induced liver injury or steatosis, (iv) Wilson disease or hemochromatosis, (v) excessive alcohol consumption (male >30 g/day, female >20 g/day), and (vi) diagnosis of malignancy within the past year. Of the eligible study participants, those with at least two of the following risk factors underwent liver biopsy: diabetes mellitus, central obesity (waist circumference ⩾90 cm for men or ⩾80 cm for women), a high level of triglyceride (⩾150 mg/dl), a low level of high-density lipoprotein (HDL)-cholesterol (<40 mg/dl for men or <50 mg/dl for women), presence of insulin resistance, hypertension, and clinically suspected MASH or fibrosis[1].

*MASLD activity scoring and fibrosis staging*

Control liver tissues were collected from subjects who underwent liver biopsy in a pre-evaluation for donor liver transplantation or in a characterization of solid liver masses that were suspected to be hepatic adenoma or focal nodular hyperplasia based on radiological results without any evidence of hepatic steatosis. All liver biopsies were assessed and reviewed by a single experienced liver pathologist. MASLD was histologically defined as the presence of ⩾5% macrovesicular steatosis with at least one of the cardiometabolic risk factors. MASH was diagnosed based on an overall pattern of histological hepatic injury consisting of macrovesicular steatosis, inflammation, or hepatocellular ballooning according to Brunt et al.'s criteria[2,3]. We also graded steatosis, lobular inflammation, and hepatocellular ballooning according to the NAFLD activity score[4]. Fibrosis was assessed according to a 5-point scale proposed by Brunt and modified by Kleiner et al.[4]: F0, absence of fibrosis; F1, perisinusoidal or periportal fibrosis; F2, perisinusoidal and portal/periportal fibrosis; F3, bridging fibrosis; and F4, cirrhosis. Advanced fibrosis was defined as F3–F4. We categorized study participants into no-MASLD control, MASL, and MASH. MASH was further classified by fibrosis severity into early MASH (eMASH; MASH with F0-2) and advanced MASH (aMASH; MASH with F3-4).

*snRNA-seq data analysis methods*

Raw reads were aligned by Cellranger[5] by using "refdata-gex-GRCh38-2020-A" downloaded from 10x genomics website as reference and "—include-introns" option. Ambient RNA was removed by Cellbender[6]. Input parameters ("—expected-cells", "—total-droplets-included") were obtained from Cellranger output file (barcodes.tsv.gz).

Output files were converted into Seurat object implement in Seurat 4[7] and further analyzed in R (version 4.1.0). Cell doublets were identified with DoubletFinder[8]. Singlet cells were filtered for gene count and read counts (count > median in each donor x 0.2). Genes expressed in less than 0.25% of cells in each donor were filtered out. Then, cells were normalized with scran (computeSumFactors)[9] and batchelor (multiBatchNorm)[10] and merged into one list. Among multiple batch correction methods, (FastMNN[10], Harmony[11], Liger[12], CCA, RPCA[7]), Liger outperformed in kBET acceptance rate[13] in our comparison analysis (Supplementary Fig. 20) and thus was utilized. Dimension reduction and clustering was performed with default Seurat methods. FindNeighbors function was used with 20 iNMF dimensions. FindClusters function was performed with varying resolutions ranging from 0.05 to 0.5, and inspected with clustree for cluster stability. Resolution of 0.2 was selected for subsequent analyses.

General cell types were annotated by inspecting cell type marker expression (Supplementary Table 2 and 12, Supplementary Fig. 2). For non-immune cells, each cell type was separately subjected to sub-cell-type annotation by marker gene expression. First, we performed subclustering and sub-cell-type-annotation among all donor cells. Next, to interrogate the difference between diseased and control cells, we subset cells from NALFD donors or control donors and performed sub-clustering separately. Finally, cells that have consistent sub-cell-annotation from both analyses were assigned to such sub-cell-type. Inconsistent cells were denoted as unidentified. Subclusters with ectopic marker gene expression were annotated as miscellaneous (misc) cell type.

Immune cells were annotated using Azimuth[7] and tested for cell type marker gene expression. Each cell-type were checked for marker gene expression and few sub-cell-types were merged or re-annotated (Supplementary Table 12). Immune cells annotated as erythrocyte (Eryth) and innate lymphoid cells (ILC) were annotated as unidentified cell type.

At the sample level, samples with low gene counts (median genes per cell < 200) were excluded. Normalized gene counts were averaged into pseudobulk expression per donor and following principal component analysis (PCA) revealed one outlier sample, which was excluded for further analysis. Three additional samples with low genome-transcriptome concordance tested by VerifyBamID[14] was excluded. Differential expression analysis was performed with Seurat FindMarkers. Intercellular interaction was analyzed by CellChat[15]. Transcription factor activity was quantified with pySCENIC[16,17]. Gene ontology enrichment was tested with enrichR[18] or ToppGene suite[19].

*hdWGCNA on snRNA-seq data*

Cells from four major cell types (hepatocytes, cholangiocytes, stellate cell, and endothelial cells) were separated into different Seurat objects, and each were subjected to co-expression network analysis using hdWGCNA package[20,21] in R. *SetupForWGCNA* function was performed with “gene_select=’fraction’, fraction=0.05” option. *MetacellsByGroups* function was used with donor and sub-cell-type as grouping variable and k=50. Then, *NormalizeMetacells, SetDatExpr, TestSoftPowers* function was used sequentially. ConstructNetwork was used with pre-selected soft power. *ScaleData, ModuleEigengenes, ModuleConnectivity* functions were consecutively used. Gene list per module was extracted by GetModules function. Harmonized module expression values were obtained by GetMEs function.

For module projection, public human liver single cell data[22-24] was downloaded and converted into SeuratObject. Functions *ProjectModules, ModuleConnectivity, SetDatExpr, ModulePreservation* were consecutively used to project hepatocyte modules and calculate preservation scores. Seurat object resulting from above hdWGCNA was used as reference, public data was input to query data.

*Genome data generation and processing*

DNA samples were extracted from the whole blood or liver tissues using ExgeneTM Blood SV mini or Exgene Tissue SV kit (GeneAll Biotechnology Co. Ltd., Seoul, Korea). DNA was shipped to Gencove (New york, NY) for 1X low-coverage WGS. Sequencing, alignment, variant calling, and imputation was performed by Gencove imputation pipeline[25] with GRCh37 v2.4 as reference. Variants were lifted over to GRCh38 using *LiftoverVcf*. One sample that failed for low coverage WGS was subjected to 30x whole genome sequencing, and variant calling was performed following this study[26], VCFs from 54 samples were merged, left-aligned with bcftools, and converted into plink[27] format. Variants were filtered through 3 steps. First, low quality and rare variants were identified separately within no-MASLD or MASLD donors (--hwe 1e-5, --geno 0.05, --maf 0.05). Overlapping variants were retained for further analysis. Second, variants located in repeat regions (union of LCR, REPEATMASKER, WINDOWMASKER_SDUST) were removed. This was because variants in such regions exhibited low genotype accuracy compared to 30x WGS in our cohort (Supplementary Fig. 21). Last, to reduce multiple testing and minimize variants with highly correlated genotypes, we clumped variants with LD R2 > 0.9 in 250kb region (--clump-r2 0.9, --clump-kb 250), and left one index variant with the highest allele frequency (Supplementary Fig. 22) for further analysis.

*Liver-eQTL and ieQTL calling: Poisson mixed effects (PME) model*

Among 48 donors, we excluded 4 donors who had less than 5 cells in any of the four major cell types (hepatocyte, cholangiocyte, stellate cell, endothelial cell; Supplementary Table 2). Poisson mixed model[28] is constructed with UMI counts as output variable, genotype and covariates as fixed effects, and donor as random effects. Covariates included disease status (binary), scaled biopsy age, sex, scaled library size (scaled nCount_RNA), five gene expression PCs. Null PME model is represented as follows:

$$\log(UMI_{c,g}) = \beta_{GT}X_{d,GT} + \beta_{MASLD}X_{d,MASLD} + \beta_{age}X_{d,age} + \beta_{sex}X_{d,sex} + \beta_{GT}X_{GT,d} + \beta_{lib\_size}log10(X_{c,lib\_size}) + \sum_{j=1}^{5}\beta_{GE\_PC_k}X_{c,GE\_PC_k} + (\phi_d|d) + \varepsilon$$

$UMI_{c,g}$ represents read count of gene g in cell c, *GT* represents number of alternative alleles of a cis-SNP in donor *d* which takes a value in [0,1,2], *MASLD* denotes binary disease status, age denotes scaled age at biopsy, $GE\_PC_k$ denotes gene expression kth PC of cell *c*. Since our cohort consists entirely of ethnic Koreans, we did not include genotype PC as covariates.

Regression model was fit with *glmer* function in lme4[29] package. Filtered and clumped variants within 1Mb distance from each gene's TSS were tested. *P*-values were adjusted for multiple testing in hierarchical manner for each cell type eQTL results. For every tested gene, *P*-values from tested *cis*-SNPs were adjusted with Benjamini-Hochberg process (Padj_SNPwise). Best Padj_SNPwise value per gene was then collected, and again adjusted with the Benjamini-Hochberg process resulting in gene-wise adjusted *P*-value (Padj_Genewise). Maximum Padj_SNPwise value that has Padj_Genewise < 0.05 was used as a *P*-value threshold. SNPs with Padj_SNPwise lower than that threshold was thought significant liver-eQTLs. Same approach was utilized to subset of cells from MASLD donors or No-MASLD donors to identify MASLD-eQTLs or control-eQTLs, correspondingly. Such eQTL calling was separately performed for hepatocyte, cholangiocyte, stellate cell and endothelial cells.

Among significant eQTLs (union of liver-eQTLs, MASLD-eQTLs, control-eQTLs), we tested for interaction with cell state represented by gene module expression or disease status of the donor. Full model with interaction term is represented as follows:

$$\log(UMI_{c,g}) = \beta_{GT}X_{d,GT} + \beta_{state}X_{c,state} + \beta_{interaction}X_{c,state}X_{d,GT} + \beta_{NAFLD}X_{d,NAFLD} + \beta_{age}X_{d,age} + \beta_{sex}X_{d,sex} + \beta_{GT}X_{GT,d} + \beta_{lib\_size}log_{10}(X_{lib\_size,c}) + \sum_{j=1}^{5}\beta_{GE_{PC_k}}X_{c,GE_{PC_k}} + (\phi_d|d) + \varepsilon$$

Cell states were defined as biologically interpretable module expression (continuous) or disease status (binary) of each cell. Module expression values were scaled to mean 0, variance 1 prior to ieQTL calculation.

Likelihood ratio test using *anova* function in R was performed to compare full model with or without interaction term. Specifically, $\chi^2$-statistic was calculated as $-2 \times \log(likelihood\ ratio)$, which was compared against $\chi^2$ distribution with one degree of freedom. P values were adjusted with the Benjamini-Hochberg method for multiple testing. ieQTLs with adjusted P value < 0.05 were considered as significant.

*Replication of liver-eQTLs*

We tested for replication of liver-eQTLs in 3 different dataset or models. First, for the pseudobulk-linear model, we averaged the single cell gene expression per cell type and donor, and applied inverse normal transformation. Genes expressed in more than 1% of cells from the cell type and 50% of donors in the cell type was only retained. Median library size per cell type x donor, age, sex, and first PEER factor was added to the linear regression model as covariates. lm() function implemented in R was applied. Hierarchical multiple testing adjustment was performed with the Benjamini and Hochberg method SNP-wise first, then Gene-wise. eQTLs with adjusted P value below 0.1 was denoted as significant.

Second, for bulk eQTLs, raw counts of 120 liver bulk RNA-seq from our previous study[30] were normalized by DESeq2, and the inverse normal transformed. Sex, age, disease status, 5 expression PCs were included as covariates. lm() function implemented in R was applied. Hierarchical multiple testing adjustment was performed as same as above, with a significance cutoff of 0.05.

For GTEx eQTLs, significant variant-gene pairs were downloaded from the website (GTEx v8).

Standardized effect sizes were calculated by beta/standard error, and gene-snp pairs that were significant in both datasets (ours and public) were used for correlation calculation.

*Stratified LD score regression (S-LDSC)*

GWAS summary statistics were converted to sumstats format. Partitioned and non-partitioned LD score was estimated from genotype data used for eQTL calling. Recombination map was downloaded from SHAPEIT5 github (https://github.com/odelaneau/shapeit5/tree/main/resources/maps/b38) and added to genotype files by plink. Heritability partitioning was performed by ldsc.py software, with gwas sumstats, partitioned LD score, univariate LD score as inputs, and "–overlap-annot, --print-cov, --print-coefficients, --print-delete-vals, --not-M-5-50" as options.

*eQTL power estimation*

Power analysis was performed following the previous study[31], assuming a linear model.

$$y_i = \beta x_i + \varepsilon_i$$

$$\varepsilon_i \sim \mathrm{N}(0, \sigma^2)$$

Here, $y_i$ and $x_i$ represents the gene expression and a specific SNP genotype of individual i (i=1..n), correspondingly. When $x_i$ are centered at 0 and scaled to variance

1, effect size $\hat{\beta}$ follows the normal distribution of mean $\beta$ and variance $\frac{\sigma^2}{n}$. Assume a standardized effect size $\lambda = \frac{\beta}{\sigma}$ which follows

$$\hat{\lambda} \sim N(\lambda, \frac{1}{n})$$

Then, power is calculated as following

$$Power = \Phi(\Phi^{-1}\left(\frac{\alpha}{2}\right) + \lambda\sqrt{n})$$

where $\alpha$ represents the significance level, set to 0.05 / 20000 = $2.5 \times 10^{-6}$ (Bonferroni's correction for 20000 tests) and $\Phi$ represents the Gaussian cumulative distribution function.

*Correlation with public snRNA-seq study*

Fully annotated human liver snRNA-seq dataset from a previous study[24] was downloaded. Our data was split into cells from No-MASLD and MASLD donors. Highly variable genes (n=2000) were calculated for each cell type data. Union of variable features from public data and our data were used for correlation calculation. Expression matrices were normalized and scaled to mean=0 and variance=1 and were input to calculate Pearson correlation coefficient. *cor.test* function implemented in R stats package was used.

*Cholangiocytes pseudotime and developmental pathways expression scoring*

Pseudotime analysis was performed with Slingshot[32], with umap coordinates as input data, clustering by resolution=0.05 as cluster labels. Starting cluster was manually selected by inspecting marker gene expression.

Genes representing progenitor or differentiated cholangiocyte states were obtained from previous studies[24,33] and documented in Supplementary Table 6. To identify a trend between pseudotime and normalized expression values, we utilized Nadaraya–Watson kernel regression. Pseudotime values were input as input x values, and normalized gene expression values were input as input y values in *ksmooth* function in R. Twenty times of bandwith determined by bw.SJ function was used to minimize the impact of gene expression noise. Fitted y values were used for visualization (Fig. 1f).

List of genes involved in Notch, Wnt, hedgehog and Hippo signaling were obtained from KEGG database[34]. The full list of genes is provided in Supplementary Table 6. Expression of each pathway was calculated with *AddModuleScore* function implemented in Seurat. Briefly, average expression of each gene is subtracted by aggregated expression of control gene sets. Control genes were randomly selected from the same expression level bin.

*Visualization of liver-eQTLs, ieQTLs*

For liver-eQTL visualization, gene counts of each cell were log2 transformed with pseudocount of 1, and cells were grouped and averaged per donor, resulting one dot per donor in the liver-eQTL plots.

For module interacting ieQTL visualization, gene counts were log2 transformed with pseudocount of 1, then cells were first grouped by donor, then grouped by module expression. Cells from one donor were equally split into three bins by module expression. For each bin, transformed counts were averaged per donor. Consequently, all module expression bins include expression values from all donors.

*Regression of EFHD1, FOXO1 expression levels*

This analysis was restricted to cells from MASLD donors, which exhibited a very strong interaction between module expression and eQTL. In addition, we performed linear regression on normalized counts from *EFHD1* or *FOXO1*. For regression analysis, normalized counts of *EFHD1* were the response variable, normalized counts of *FOXO1* were the explanatory variable, and expression PC_1, sex, biopsy age were input as covariates in a linear regression model.

*In vitro translation of FOXO1*

FOXO1 proteins were in vitro translated using the TnT T7 Quick Coupled Transcription/Translation System (Promega, Madison, WI, USA). 40ul of Quick Mastermix, 400ng of pcDNA3 Flag FKHR vector (Addgene #13507), 1ul of 1mM Methionine with a total volume of 50ul was mixed and incubated at 30’C for 80 minutes. Protein lysates were aliquot into 10ul and stored at -80’C before use.

*EMSA*

Electrophoretic Mobility Shift Assays (EMSAs) were performed using the LightShift Chemiluminescent EMSA Kit (Thermo Fisher Scientific, Waltham, MA) following a previous study[35]. Unlabeled and 3' biotinylated single-strand oligonucleotide probes were obtained from Bioneer (Daejeon, South Korea) and annealed by heating to 95°C followed by gradual cooling. The binding reaction consisted of 2 $\mu$L of 10X binding buffer, 500 ng of Poly(dI:dC), 400 fmol of biotin-labeled probe, and 4 $\mu$L of in vitro translated FOXO1 lysate, totaling 20 $\mu$L. For competition assays, a 10X molar excess of unlabeled probe was included. Binding reactions were pre-incubated for 10 minutes before adding the biotin-labeled probe, then incubated at room temperature for 90 minutes. 5ul of 5X loading dye was added, and 20ul of samples were electrophoresed on a 6% polyacrylamide gel at 100V in 0.5X TBE at 4°C for 55 minutes. The gel was then transferred to a positively charged nylon membrane (Roche, Basel, Switzerland) at 4°C for 30 minutes at 380mA in 0.5X TBE. DNA was cross-linked using UV light for 15 minutes with the EtBr program on a ChemiDoc MP imaging system (Bio-Rad, Hercules, CA). Biotinylated probes were detected using the chemiluminescent nucleic acid detection module (Thermo Fisher Scientific, Waltham, MA, USA), and images were captured with the ChemiDoc MP imaging system. Probe sequences are available in Supplementary Table 13. Experiment was performed 3 times.

*Luciferase reporter assay*

Reference genome sequences of 501 bp length, containing reference or non-reference eSNP allele at the mid-point were used as enhancer region (chr2:232655294-

232655794, GRCh38). Enhancer region was cloned into the upstream of luciferase gene in pGL3 promoter backbone by Vectorbuilder (Chicago, IL; Supplementary Table 14).

HepG2 cells were cultured in DMEM high glucose with 10% FBS and penicillin-streptomycin, and seeded at 2 x $10^4$ cells per well in 96 well plate. Twenty-four hours later, 10:1 admixture of enhancer luciferase vector with Renilla control vector was transfected into cells with lipofectamine 3000 (Invitrogen, Waltham, MA) according to the manufacturer's protocol. On the following day, cells were treated with 1 uM FOXO1 inhibitor (AS1842856, Sigma, St. Louis, MO) or same volume of DMSO as control. Twenty-four hours later, cells were harvested and dual luciferase reporter assay (Promega, Madison, WI) was performed in accordance with the manufacturer's instructions. Experiments were repeated 3 times with 6-7 wells per vector each time.

*Western blot validation of in vitro translated FOXO1 protein*

5ul of In vitro translated proteins was diluted with 14ul of TBS buffer and treated with 1ul of 1/10 diluted RNaseA at room temperature for 5 minutes. 5X sodium dodecyl sulfate-polyacrylamide (SDS-polyacrylamide) sample buffer was added, heated to 70'C for 10 minutes. 10ul of the prepared sample was loaded to 8% SDS-PAGE gel. Electrophoresis was performed with 80V for 15 minutes for stacking, and 120V until dye reached few centimeters above the bottom. Proteins were transferred to PDVF membrane in the transfer buffer at 160 mA for 1 hour in 4'C. Membrane was blocked with 5% skim milk for 1 hour at 4'C. Primary antibody for FOXO1 (C29H4, #2880; Cell

Signaling Technology, MA, USA) was diluted to 1:1000 and incubated with the membrane overnight in 4’C. Membrane was washed with TBST for 5 minutes, 3 times. Secondary antibody diluted in 5% skim milk was added and incubated for 1 hour. Membrane was washed, treated ECL reagents (West save star, Bologna, Italy), and detected with Chemidoc MP imaging systems (Bio-Rad, Hercules, CA).

*Generation of Efhd1-overexpressing cells*

To establish Efhd1 overexpressing cells, pCMV-SPORT6-Efhd1 plasmids were transfected to AML12 cells using Lipofector-EZ (Aptabio Therapeutics Inc., Gyeonggi-Do, South Korea) following the manufacturer’s instructions. The clone was provided from Korea Human Gene Bank, Medical Genomics Research center, KRIBB, Korea. Overexpression was validated with real time quantitative PCR (RT-qPCR, Supplementary Fig. 23).

*Western blot analysis of Efhd1-overepxressing cells*

Cell lysates were prepared using a lysis buffer containing 10 mM Tris, 100 mM NaCl, 1 mM EGTA, 10% glycerol, 1% TritonX-100, and 30 mM sodium pyrophosphate. After centrifugation at 13,000 ×g for 15 minutes, the soluble fractions of the lysates were collected. Protein samples were then separated using SDS-PAGE and transferred to nitrocellulose membranes (GE Healthcare, Madison, WI, USA). The membranes were blocked for 1 hour in 5% skim milk in phosphate-buffered saline containing 0.05% Tween-20 (PBST). Following blocking, the membranes were incubated with primary antibodies at 4 °C overnight. After washing with PBST, the membranes were incubated with horseradish peroxidase (HRP)-conjugated secondary antibodies (Cell Signaling Technology, Beverly, MA, USA) at room temperature for 2 hours. Protein bands were visualized using the Immobilon Western Chemiluminescent HRP Substrate (Merck Millipore, Billerica, MA, USA) or enhanced chemiluminescence (ECL) system reagent.

*Antibodies*

Antibodies recognizing FAS (610962) were purchased from BD Biosciences (San Jose, CA, USA). Acetyl-CoA carboxylase (3662S), SCD1 (2438S), phospho-AMPKα (2535S), Caspase 3 (9662S), cleaved Caspase-3 (9664S) antibodies were purchased from Cell Signaling Technology (Danvers, MA, USA). Anti-GAPDH (CB1001) antibody was supplied from Merck (Burlington, MA, USA). The GRP78 antibody (ab21685) was obtained from Abcam (Cambridge, UK). Anti-FOXO1 antibody (C29H4, #2880) was purchased from Cell Signaling Technology (Danvers, MA, USA)

*Measurement of Mitochondrial Membrane Potential (MMP)*

AML12 cells were treated with 2 µg/mL JC-1 (Invitrogen, MA, USA) and incubated for 20 minutes at 37 °C. Following incubation, the cells were washed twice with Dulbecco's PBS. The fluorescence signal was then measured using the IncuCyte ZOOM Live Cell Analysis System (Essen Bioscience, MI, USA).

*Real-time monitoring of cell viability using IncuCyte®*

AML12 cells ($3 \times 10^3$) were plated in a 96-well plate. Following an overnight incubation, the cells were transfected to induce Efhd1 overexpression. Twenty-four hours post-transfection, the cells were exposed to 200 ng/mL tunicamycin. Cell confluence was monitored every 4 hours using the IncuCyte® S3 Live Cell Analysis System (Essen Bioscience, MI, USA).

*Liver organoids culture*

Human liver organoids were generated from iPSCs following the previous study[36,37]. Liver organoids were cultured in Hepatic medium (HM) for one week with medium changes every 2-3 days. Enlarged organoids were mechanically divided with surgical blade and then passaged weekly at 1:3-1:5. For further differentiation of organoids, split organoids were incubated in expansion medium (EM) supplemented with 25 ng/ml BMP7 (PeproTech, Cranbury, NJ, USA) for 3 days and then in differentiation medium differentiation medium (DM) for an additional 6 days. The liver organoids are then cultured in Wiliam's E (WE) medium for additional 7 days. Detailed medium composition is described in Supplementary Table 15.

*siRNA transfection and Hepatic steatosis modeling*

Following 14 days of organoids culture, liver organoids were transfected with siRNA. Liver organoid received either 100 nM EFHD1 or non-targeting control siRNA (unlabelled). siRNA complexes were formed using RNAiMAX (Invitrogen, Waltham, MA, USA), in DMEM (Gibco, Waltham, MA, USA) containing 10% normal FBS (Gibco, Waltham, MA, USA). 350 µl of formed siRNA complex medium was then bathed over matrigel dome contained within a single well of a 24-well plate overnight. The next day, liver organoids were treated with either normal medium or high-fat medium. The high-fat medium contained 0.2 mM oleate (cat # O7501; Sigma, St. Louis, MO, USA) and 0.1 mM palmitate (cat# P9767; Sigma, St. Louis, MO, USA) complexed with 12% fatty acid-free BSA (cat# A8806; Sigma, St. Louis, MO, USA) and was treated for 2 days.

*RNA extraction and real-time polymerase chain reaction (PCR)*

Total RNA was extracted using easy-BLUE™ reagent (iNtRON, Gyeonggi-do, South Korea) according to the manufacturer's instructions. Complementary DNA was synthesized using TOPscript™ RT DryMIX (Enzynomics, Daejon, South Korea) and quantitative PCR was performed using Fast SYBR® Green Master Mix (Applied Biosystems, Waltham, MA, USA) with gene-specific primers (Supplementary Table 16). *ACTB* was used as an internal control. For every iPSC line and condition, two organoids were generated, totalling 6 organoids per genotype, condition. Two-sided t-test was performed to test for statistical significance. *P*-value below 0.05 was indicated as significant.

*Lipid droplet staining*

Organoids were washed with cold PBS (Welgene, Gyeongsangbuk-do, South Korea) and fixed with 4% paraformaldehyde (PFA) (Biosesang, Gyeonggi-do, South Korea) for 15 minutes at RT. PFA-fixed organoids were washed with PBS, and then incubated with 30% sucrose (Sigma, St. Louis, MO, USA) solution at 4 °C overnight. Dehydrated organoids were embedded in O.C.T. compound (Sakura Finetek, Torrance, CA, USA) and snap-frozen. Sections were obtained with a cryostat (LEICA, Wetzlar, Germany) as 10 µm thick. The sectioned organoids were stained with 2 µM BODIPY (Invitrogen, Waltham, MA, USA) staining solution (Thermo Fisher, Waltham, MA, USA) with 2 µM Hoechst 33342 (Invitrogen, Waltham, MA, USA) for 1 hour at RT in the dark. Images were obtained using an EVOS FL Auto 2 (Invitrogen, Waltham, MA, USA) microscopy. Images were loaded and quantified using the EBImage library in R. For the thresh() function, a half window width (parameters w and h) of 5 and thresholding offset (parameter offset) of 0.1 were used. Due to low signal intensity and background noise in two images from organoid CW10206, and three images from from CW20009, the offset was adjusted to 0.005 for DAPI image (blue) and 0.04 or 0.02 for BODIPY image (green) based on manual inspection.

### Supplementary Figures

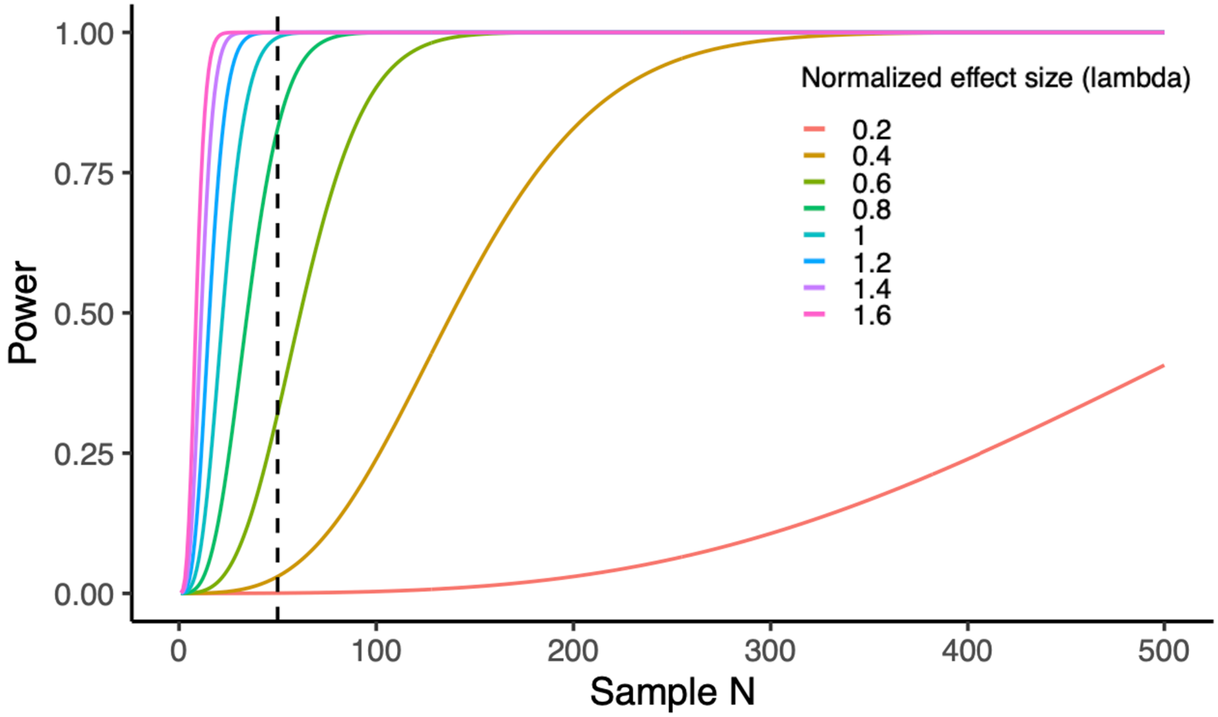


**Supplementary Figure 1. eQTL power analysis**

Power to detect eQTLs is plotted against sample size (x-axis) and standardized effect sizes (color) using a linear model. The significance threshold was set at $2.5 \times 10^{-6}$ (Bonferroni correction for 20,000 tests), with a dashed line marking a sample size of 50.

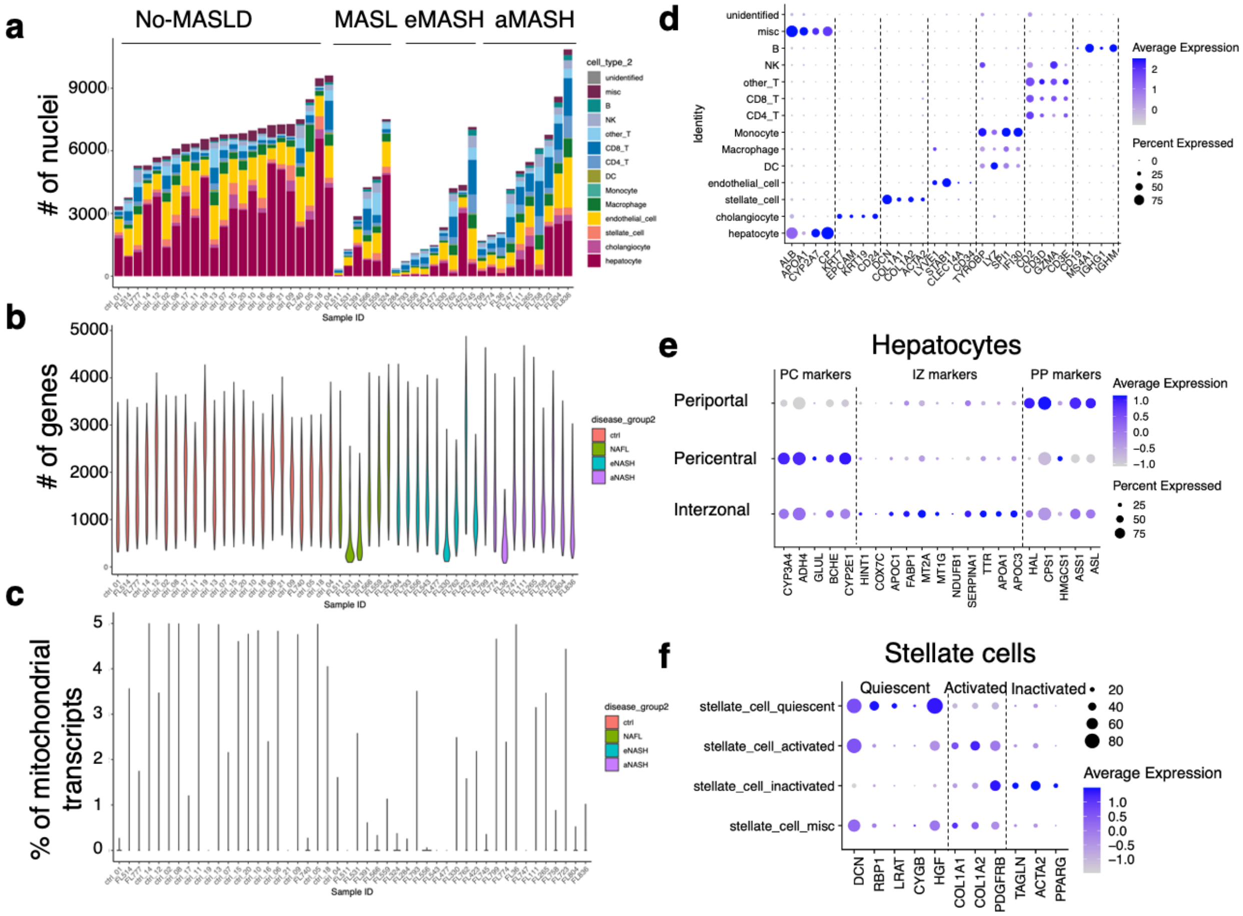


**Supplementary Figure 2. snRNA-seq data quality control metrics**

**a**, Stacked bar plot of nuclei count per sample (x-axis) and cell type (color). **b-c**, Violin plot of gene count per cell (nFeature_count) and mitochondrial transcript percent per sample, respectively. Color represents disease stage. **d**, Dot plot showing expression of marker genes across cell types. **e**, Epression of zonation marker genes in hepatocyte subclusters. **f**, Expression of stellate cell activation marker genes in stellate cell subclusters.

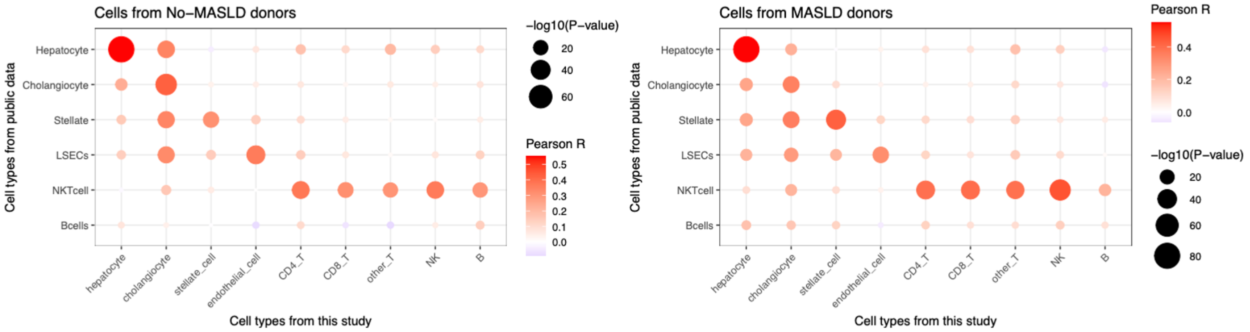


**Supplementary Figure 3. Correlation with a public snRNA-seq study**

Pearson's correlation coefficient values between single nucleus transcriptomes from our data and public data[24] are plotted. Analysis used the scaled expression values on the union set of 2,000 highly variable genes for each cell type and dataset. Cells were split by the donor's disease status (left: No-MASLD, right: MASLD). Dot size corresponds to the –log10(*P*-value), and the color represents the pearson's correlation coefficient.

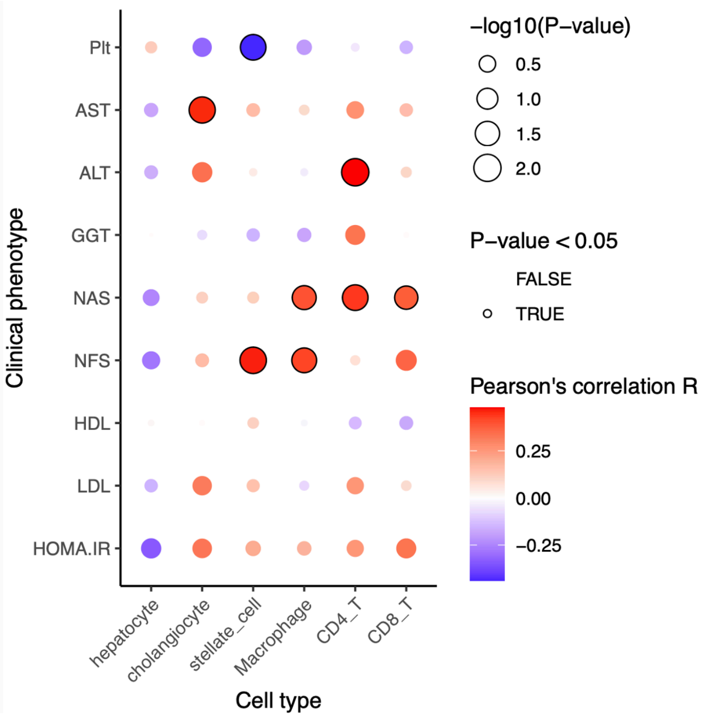


**Supplementary Figure 4. Correlation between cell type proportion and clinical features of patients**

Dot plot correlating cell type proportions per donor with various clinical phenotypes, where dot size, outline, and color represent the –log10(*P*-value), statistical significance, and the correlation coefficient, respectively.

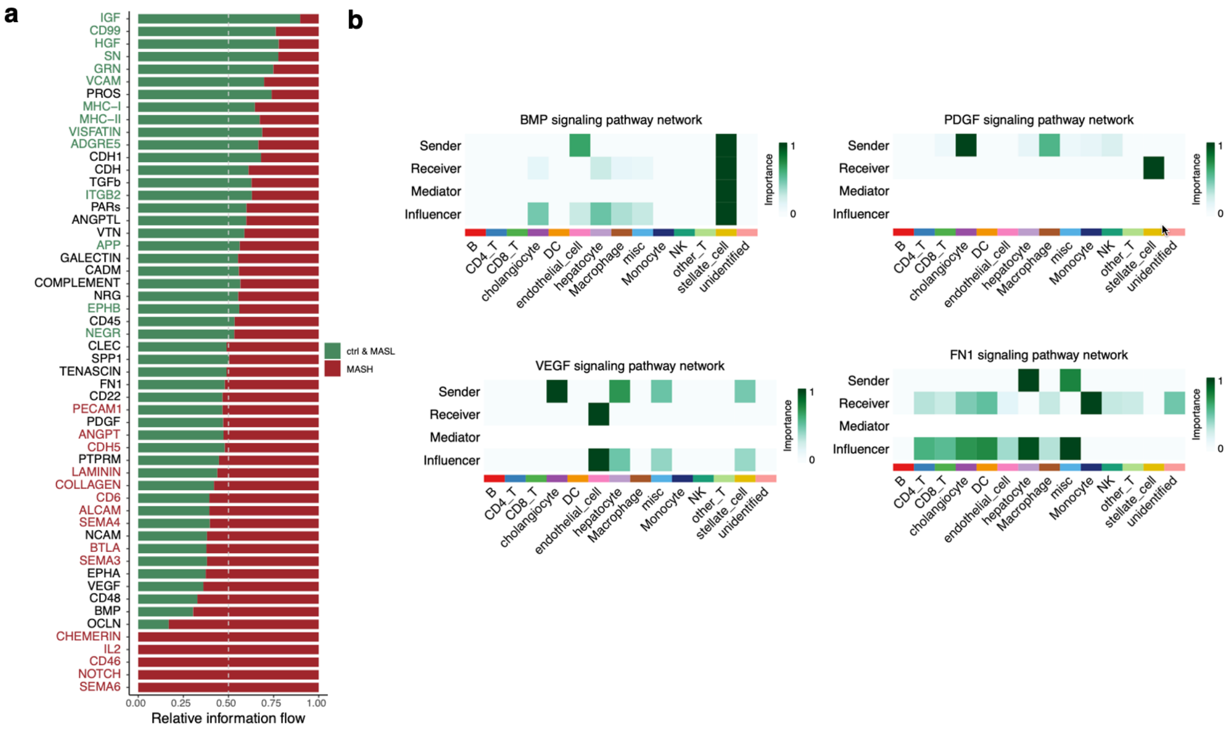


**Supplementary Figure 5. Differences in the overall cell-cell interaction patterns in No-MASH versus MASH cells**

**a**, Plot of relative information flow in various signaling pathways. Comparing No-MASH (green) and MASH (red) cells. Information flow is the sum of communication probability among all pairs of cell groups. Signaling pathways were ranked by differences in information flow, highlighted in corresponding colored text for significant Wilcoxon test results. **b**, Heatmaps of centrality scores for well-known liver signaling pathways across all donors.

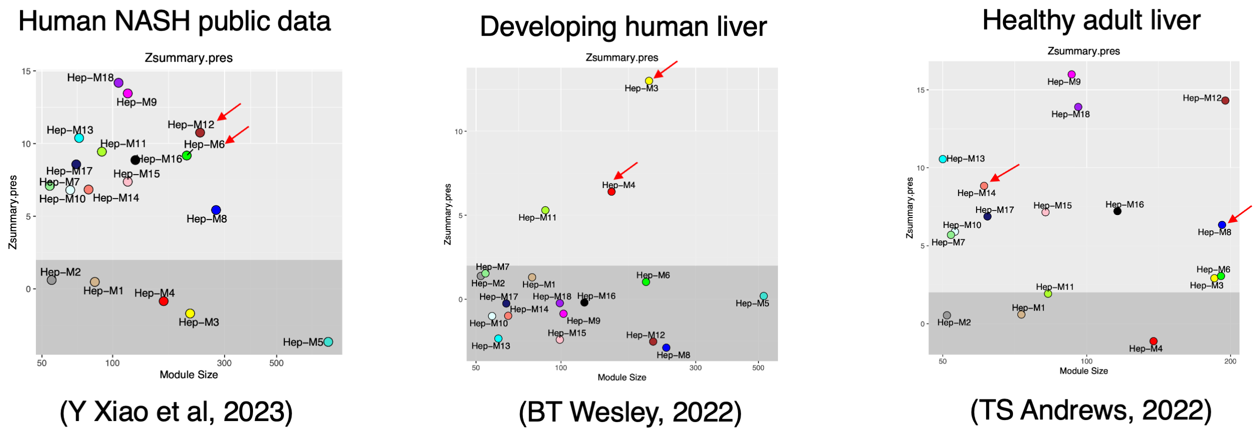


**Supplementary Figure 6. Reproducibility of hdWGCNA module networks in public data**

Scatter plot of Z-scores for module reproducibility (Zsummary.pres; y-axis) against the number of genes in the module (module size; x-axis). Reproducibility was calculated by ModulePreservation function in hdWGCNA library. Z-score exceeding 2 indicated as light gray box, indicating that the module is reproducible. Red arrow indicates the modules used for plotting Fig. 1d.

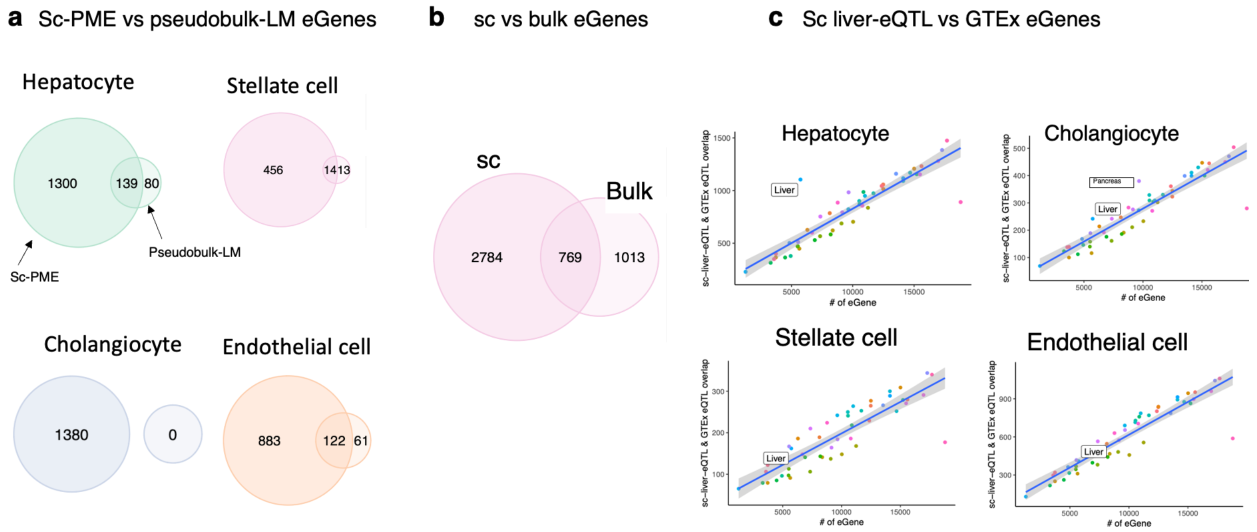


**Supplementary Figure 7. Replication of liver-eQTLs with other model or cohorts.**

**a-b**, Venn diagram of eGenes discovered by PME model and **a**, pseudobulk linear model or **b**, bulk eQTL from a different liver cohort. **c**, Number of eGenes that are replicated in GTEx eQTLs from various tissues, plotted against the number of significant eGenes in each GTEx tissue.

Abbreviations: sc-PME, liver-eQTLs called at single-cell resolution using Poisson mixed effects model; pseudobulk-LM, liver-eQTLs called from snRNA-seq data by aggregating into pseudobulk matrix and appling linear model; sc, single-cell liver-eQTLs; Bulk, bulk eQTLs called from a separate liver cohort.

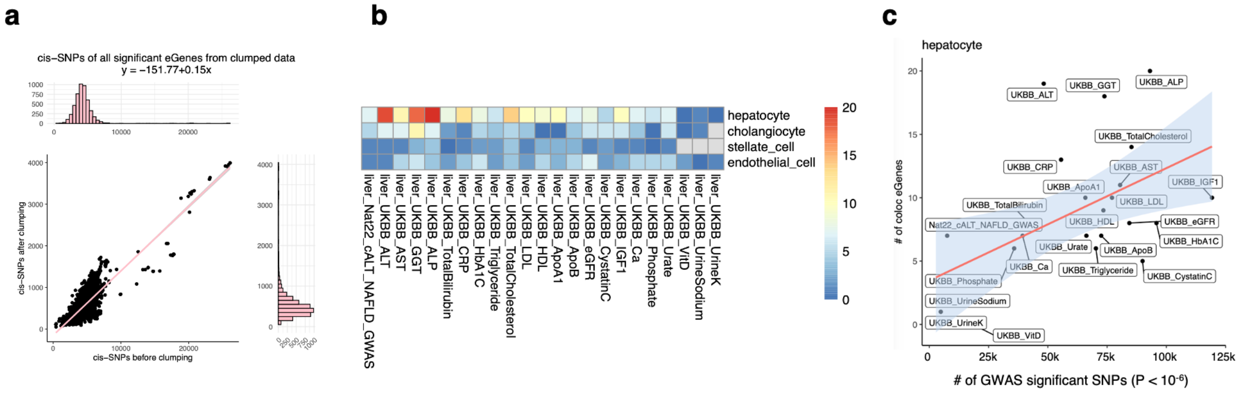


**Supplementary Figure 8. Colocalization analysis without LD clumping on cis-SNPs**

**a**, Number of cis-SNPs of significant liver-eGenes before (x-axis) and after (y-axis) LD clumping shown as scatter plot and a pink linear trend line. Histogram of each axis data is displayed parallel to the axis. **b-c**, Analogous to Fig. 2e-f, but all SNPs within 1Mb from the transcription start site of significant liver-eGene was used to calculate colocalization probabilities. **b**, Heatmap depiciting the number of liver-eGenes that colocalized with various GWAS variants, before LD clumping. **c**, Scatter plot displaying the relationship of the number of colocalizing hepatocyte liver-eQTLs (y-axis) with the number of significant GWAS SNPs from UK biobank phenotypes (x-axis). The line of best fit is shown, with the 95% confidence interval in gray.

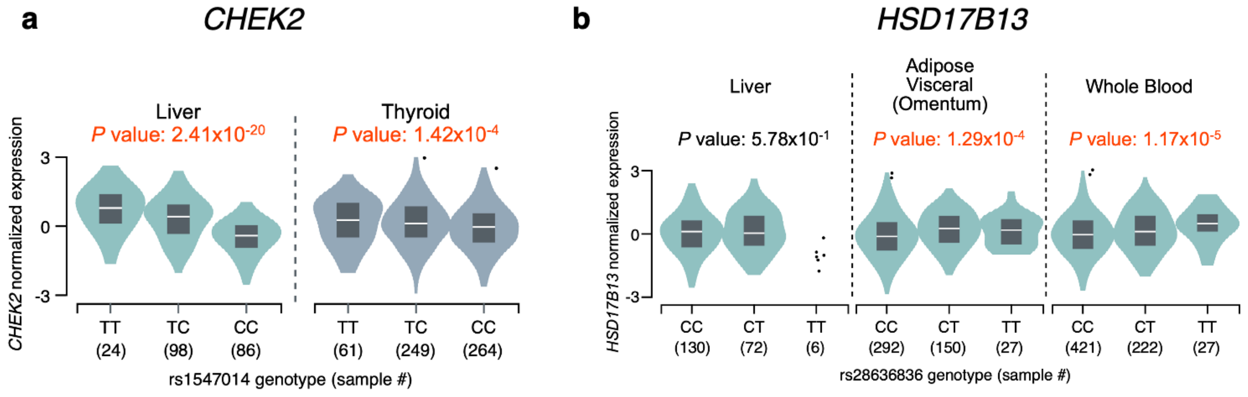


**Supplementary Figure 9. Sc-eQTLs in GTEx dataset**

**a**, eQTL plot displaying the *CHEK2* expression against rs1547014 genotype in GTEx liver and thyroid tissue. Significant eQTL *P*-values are denoted in orange on top. This eQTL was not significant in other tissues. **b**, eQTL plot displaying the *HSD17B13* expression against rs28636836 genotype in GTEx liver, skin, and whole blood. *P*-values are denoted as orange. This eQTL was also significant in 30 other tissues, but not in the liver.

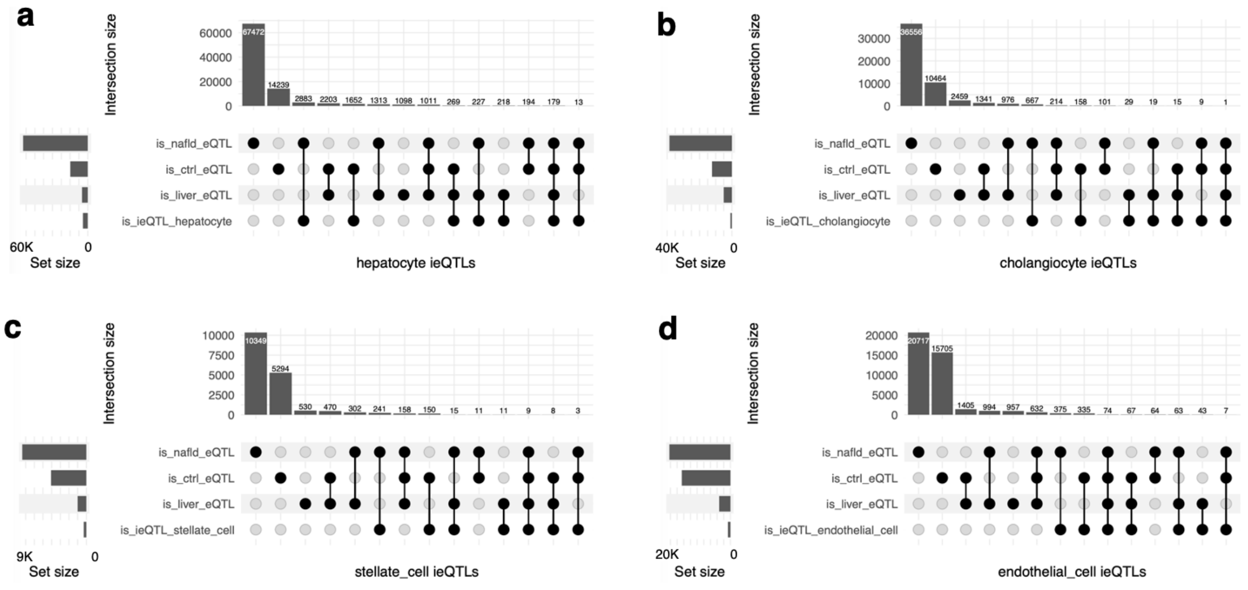


**Supplementary Figure 10. Counts of liver-eQTLs, MASLD-eQTLs, control-eQTLs and ieQTLs**

Upset plots showing the intersections among liver-eQTLs, MASLD-eQTLs, control-eQTLs, and  ieQTLs in **a**, hepatocytes, **b**, cholangiocytes, **c**, stellate cells and **d**, endothelial cells.

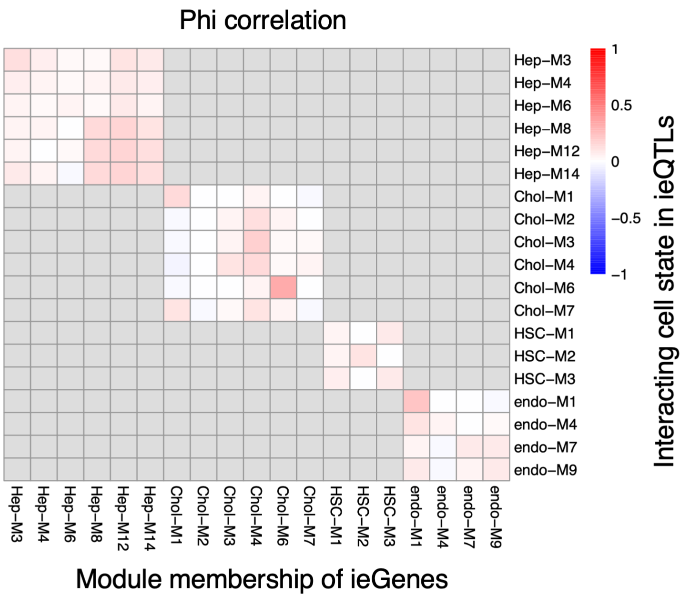


**Supplementary Figure 11. Module membership of ieGenes**

For each module, we tested how much overlap is between 1) the genes assigned to that module (x-axis), and 2) ieGenes that show significant interaction with the cell state (y-axis). The phi correlation coefficient was calculated and displayed as a heatmap.

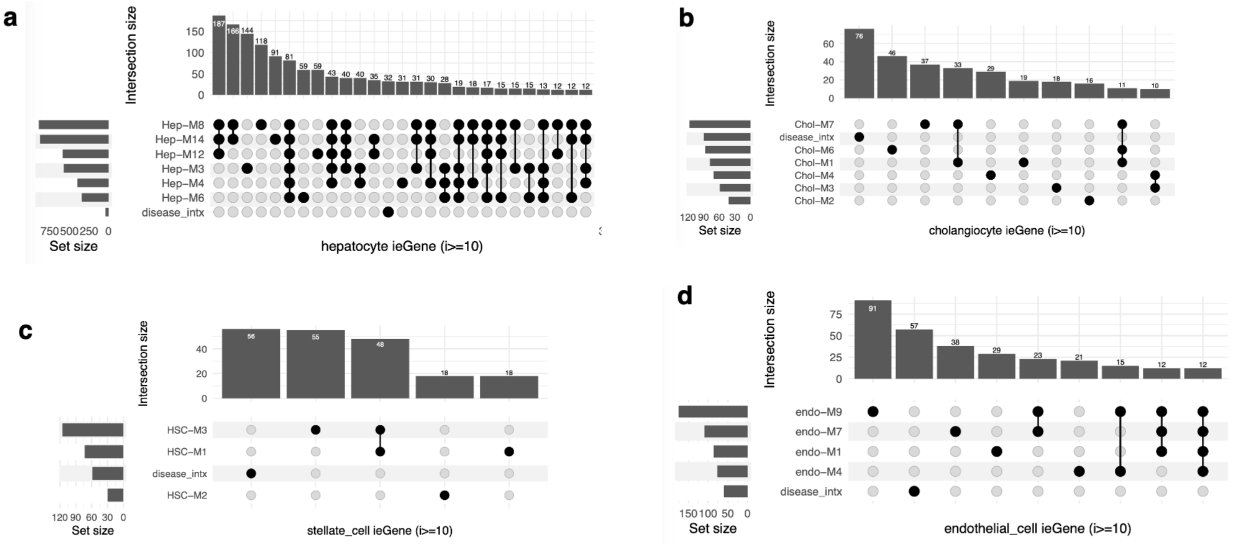


**Supplementary Figure 12. Upset plot of cell states that had significant ieQTLs**

Upset plot of ieGenes that interact with different cell states (module expression) in **a**, hepatocytes, **b**, cholangiocytes, **c**, hepatic stellate cells, and **d**, endothelial cells.

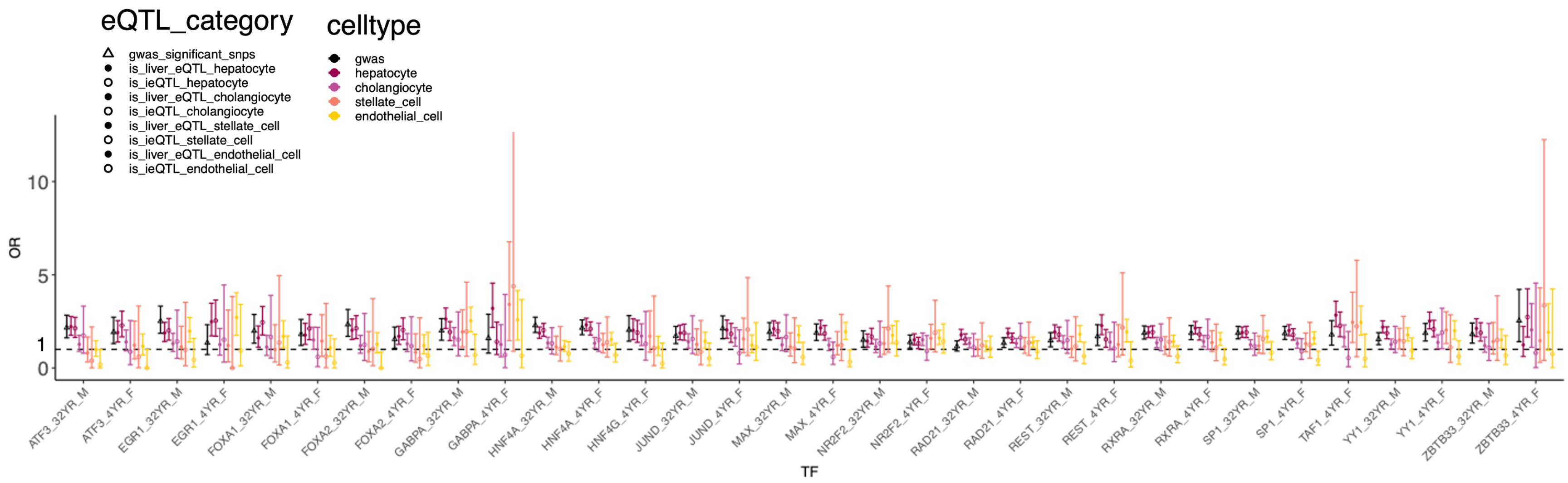


**Supplementary Figure 13. eSNPs are enriched in known TF binding sites**

liver-eQTLs and ieQTLs were annotated with ENCODE transcription factor (TF) ChIP-seq peaks, and tested for enrichment compared to all other SNPs, using fisher's exact test. Odds ratio and 95% confidence interval are plotted. X-axis represents the ENCODE ChIP-seq dataset, coded as TF_[age of the donor]_[sex of the donor]. Y-axis denotes the odds ratio. Horizontal dashed line marks an odds ratio of 1.

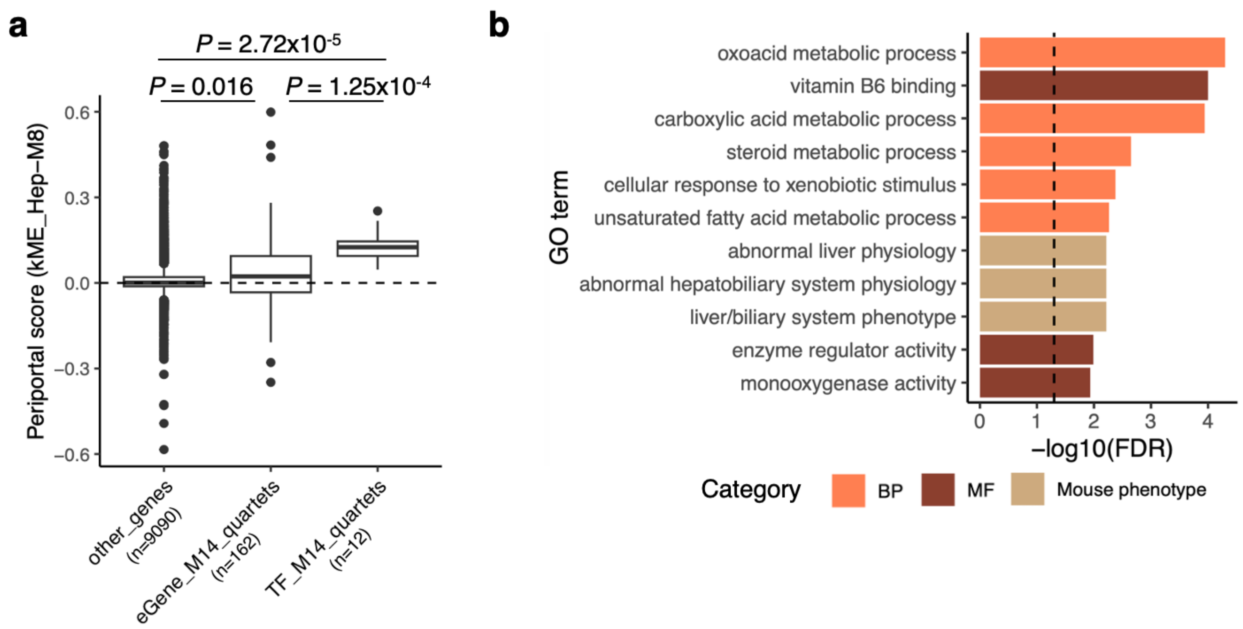


**Supplementary Figure 14. Analysis of quartets associated with Hep-M14 or M12 module-related cell states. a**, Boxplot showing the periportal-biased expression of eGenes and TFs included in quartets associated with the Hep-M14 module (periportal zone-associated). The y-axis represents the periportal score, calculated as the Pearson correlation coefficient between the gene's expression profile and the Hep-M14 module eigengene (*i.e.*, kME). t-test *P* values are indicated at the top. 'Other_genes' refers to hepatocyte-expressed genes not classified as eGenes or TFs within the M14-associated quartets. The dashed line indicates y=0. **b**, GO terms enriched in eGenes from Hep-M12 (healthy hepatocyte signature) associated quartets, highlighting metabolic pathways prevalent in the liver. The dashed vertical line indicates FDR=0.05. Abbreviation: FDR, false discovery rate

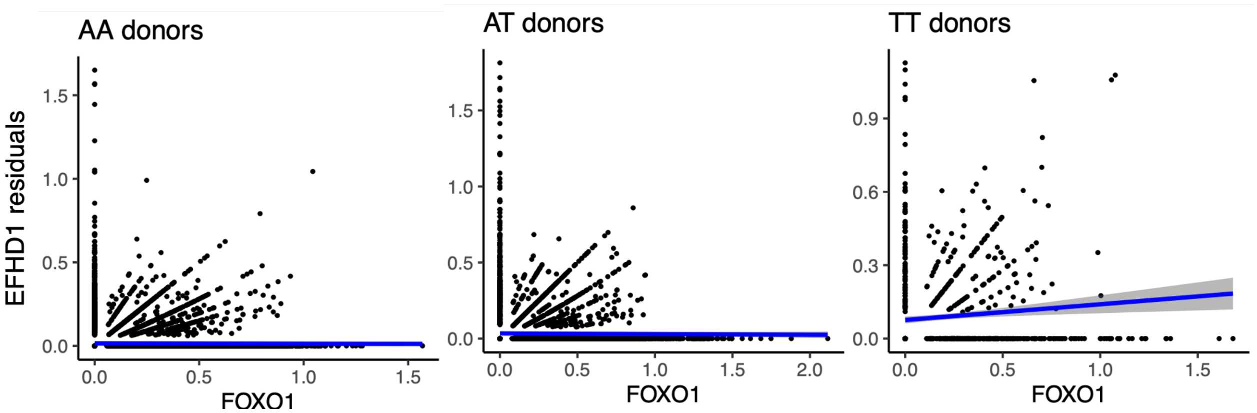


**Supplementary Figure 15. Correlation between *FOXO1* and *EFHD1* expression**

Cells from MASLD donors were split by rs13395911 genotype, and correlation between *EFHD1* expression and *FOXO1* expression are plotted. Covariates (Expression PC1, sex, and age) were regressed out from *EFHD1* expression, and its residuals were plotted against *FOXO1* expression. Line of best fit and its confidence interval are plotted as blue line and gray area, respectively. Only cells from TT donors (right) showed significant correlation ($P = 0.01$).

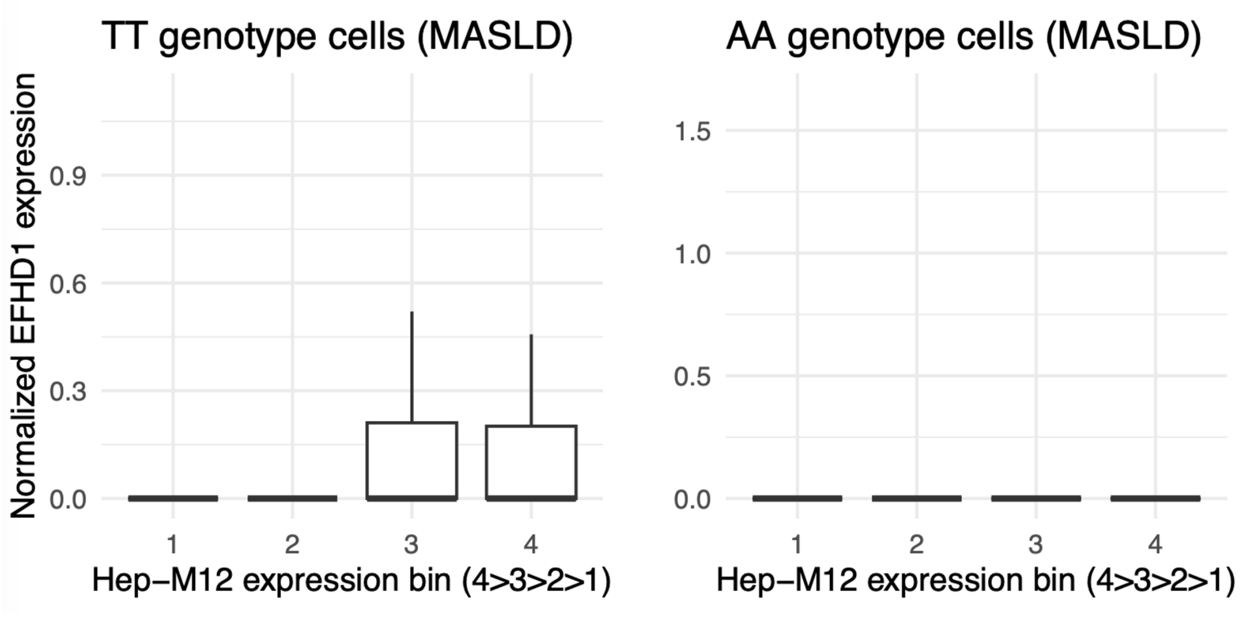


**Supplementary Figure 16.** ***EFHD1*** **expression fold change in different Hep-M12 quartiles in AA/TT donors**

Cells from MASLD donors were grouped by genotype, and sorted into four bins by Hep-M12 module expression level. *EFHD1* expression was averaged among cells from each bins and displayed into a boxplot.

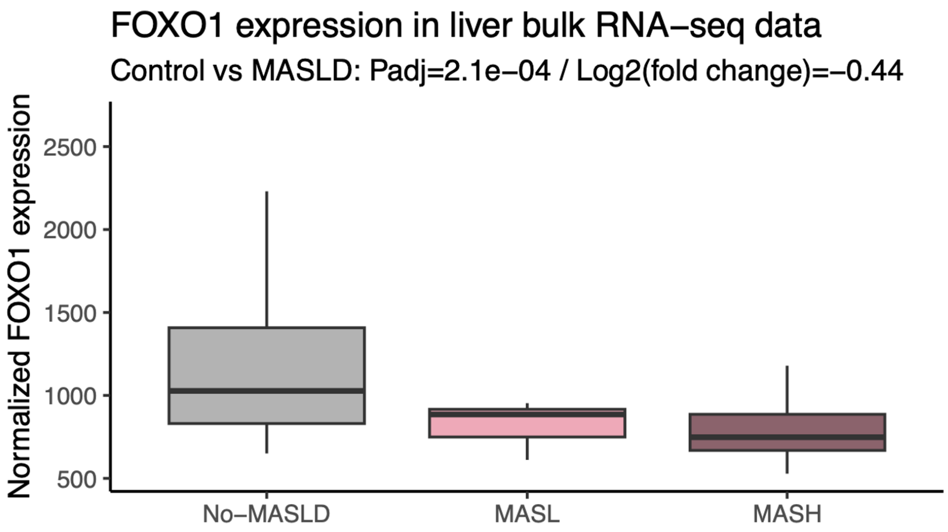


**Supplementary Figure 17. Loss of *FOXO1* expression during MASLD progression**

Bulk RNA-seq data on biopsied liver tissue from human MASLD cohort was obtained from a previous study[30] to calculate *FOXO1* expression against the disease stage. R library DESeq2 was used to normalize counts and calculate differential expression statistics.

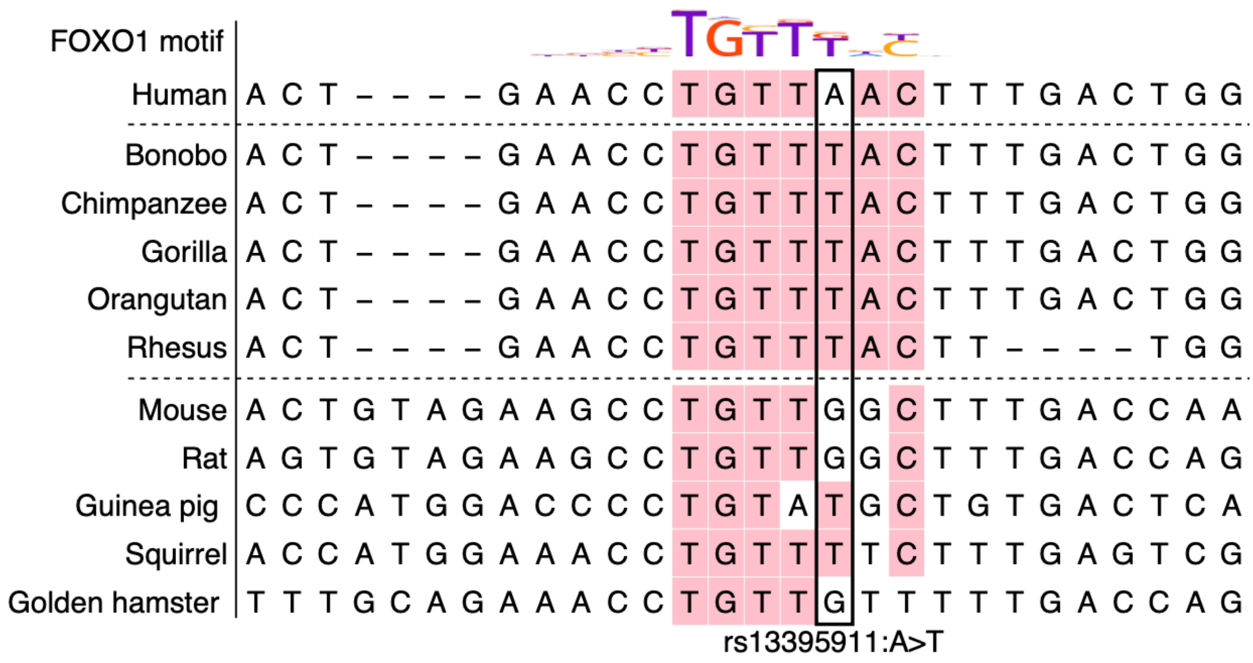


**Supplementary Figure 18. Multiple alignments of sequences near rs13395911 in human, primates and rodents.**

FOXO1 motif is shown as a sequence logo at the top. Sequences matching the FOXO1 motif are colored in pink. The position of rs13395911 is marked with a black box.

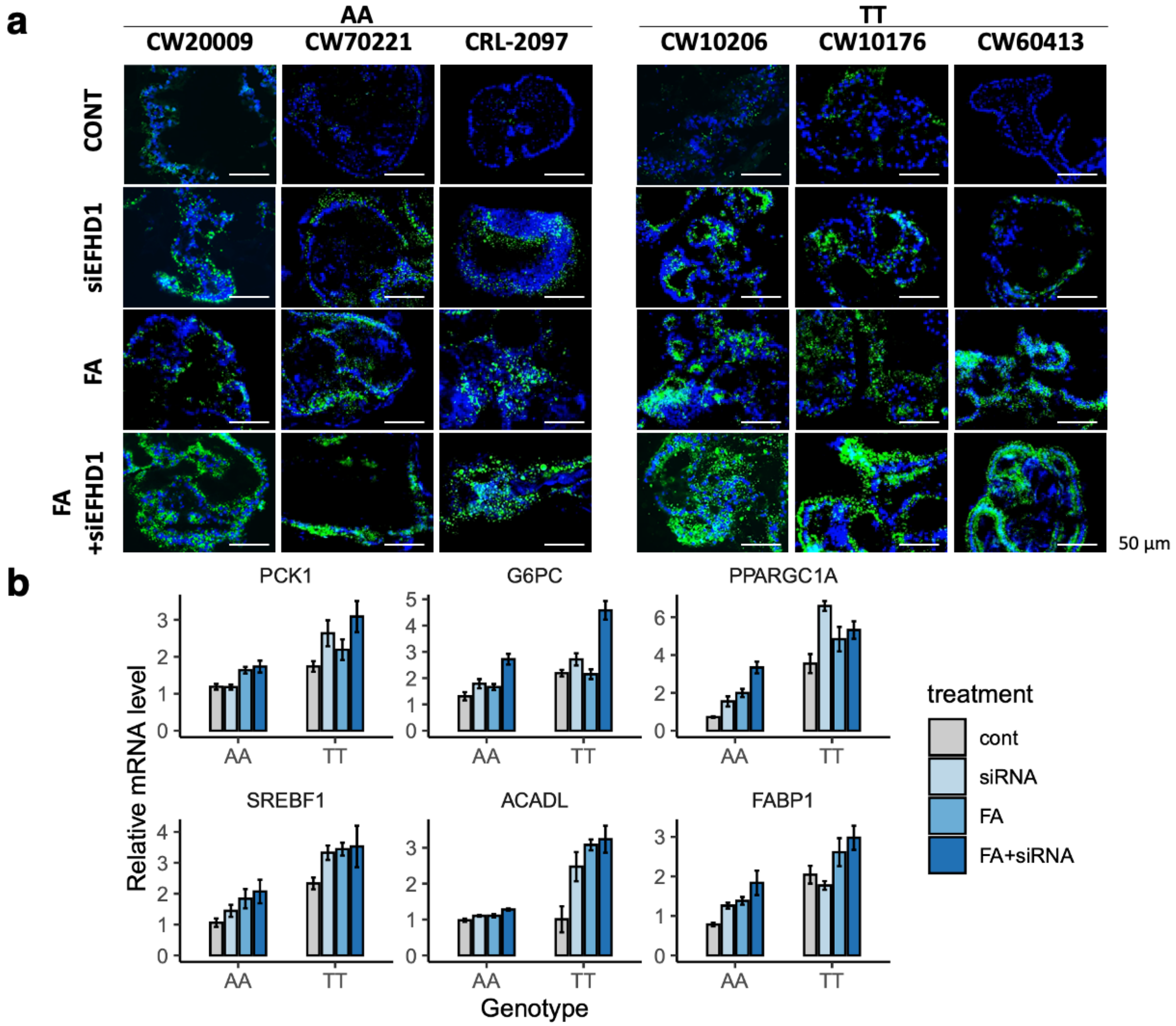


**Supplementary Figure 19. Lipid staining and RT-qPCR results from human hepatic organoids.**

**a**, Microscopic images of lipid droplet-stained hepatic organoids. Green fluorescence is from BODIPY, staining the lipid droplets. Blue fluorescence is from Hoest33342, staining the nucleus. Name of iPSC lines and their rs13395911 genotype are denoted on top. Treatment conditions are denoted on left. **b**, RT-qPCR results from hepatic organoids. Mean values with error bars representing standard error are shown.

Abbreviations: cont, control; FA, free fatty acid treatment

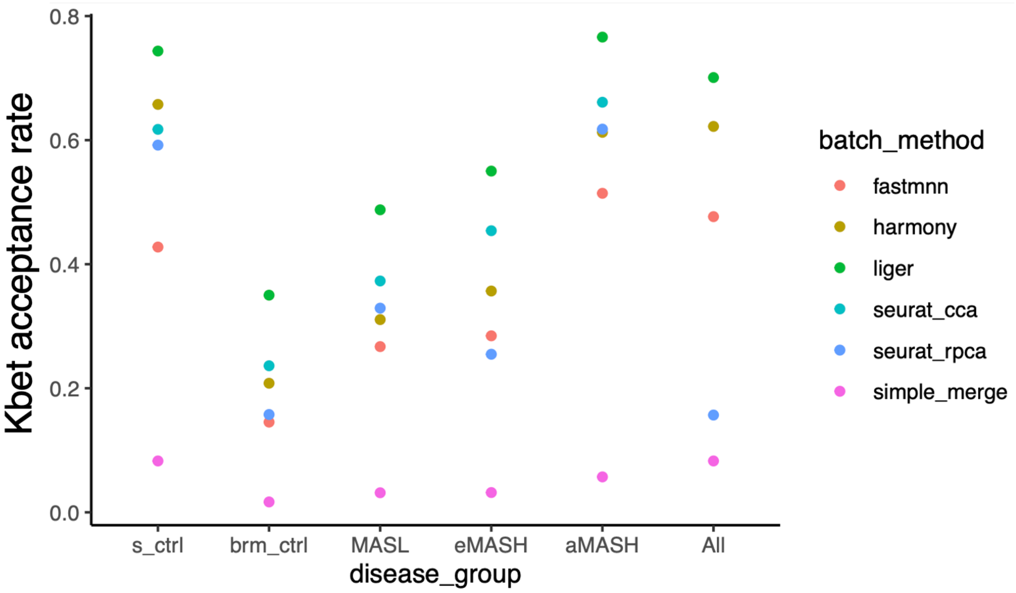


**Supplementary Figure 20. Comparison of various batch correction methods**

Evaluation of batch effect correction methods by k-bet acceptance rate in cells in subgroup of cells. S_ctrl and brm_ctrl are both control samples but split by hospital of sample origin. Five batch correction algorithms were applied for k-bet calculation. The Liger method (represented as green), which showed the highest k-bet acceptance rate, was used for further analysis.

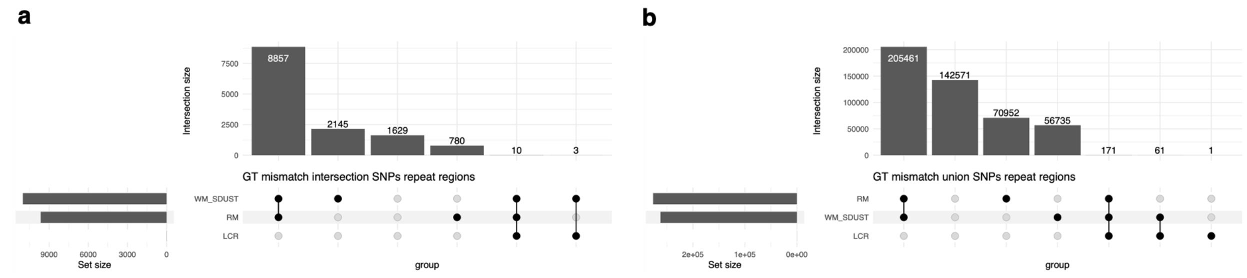


**Supplementary Figure 21. Genotyping accuracy in repeat regions**

Variants that had different genotype calls from 30X WGS and low-coverage WGS (i.e. GT mismatch variants) were annotated for various repeat regions, and their counts are plotted into upset plots. Among GT mismatch variants from three samples in which we performed both 30X WGS and low-coverage WGS, we used an intersection set in **a** or union set in **b**.

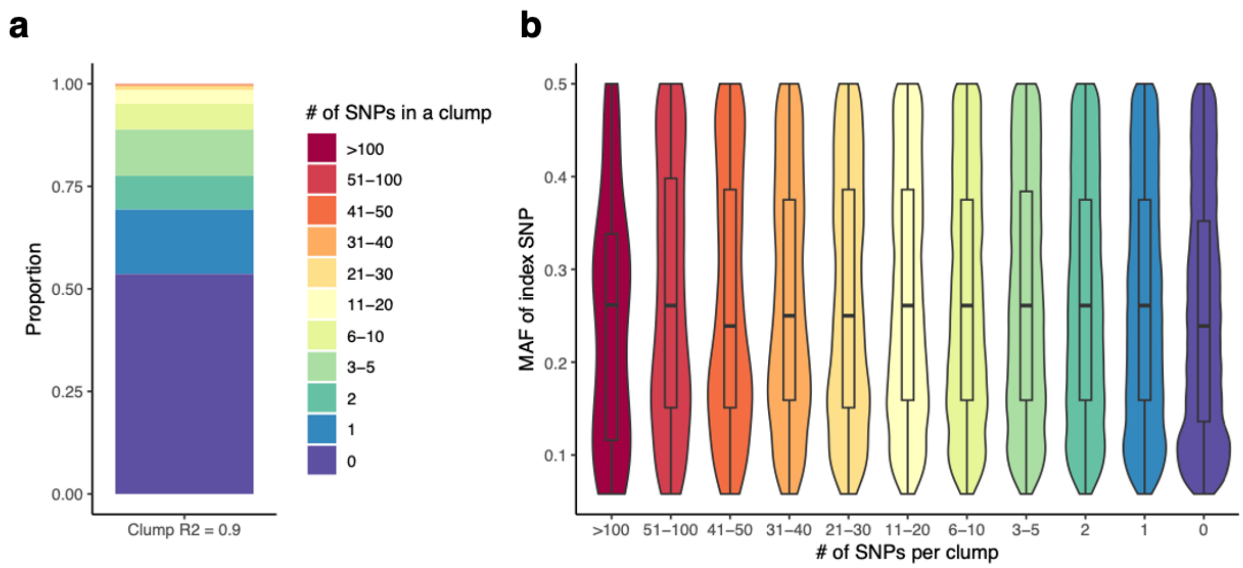


**Supplementary Figure 22. Distribution and size of LD clumps**

Prior to eQTL calling, SNPs within $R^2$>0.9 were clumped together, and an index SNP with the highest minor allele frequency was retained. **a**, Stacked bar plot showing the distribution of clump sizes, with color representing the number of SNPs excluding the index SNP within each clump. **b**, Box and violin plot depicting the minor allele frequency of the index SNP relative to the clump size.

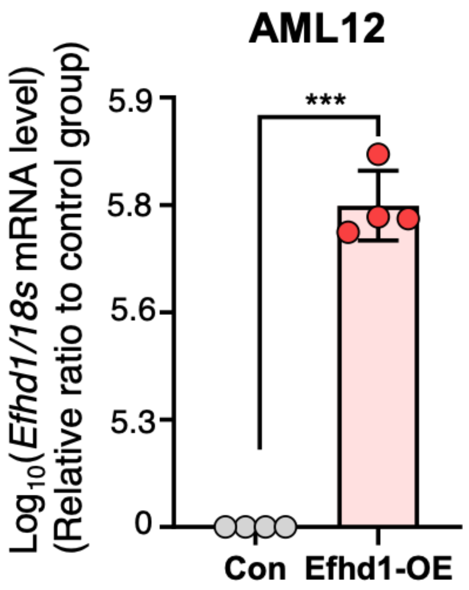


**Supplementary Figure 23. Overexpression of Efhd1 in AML12 cells validated with RT-qPCR.**

Relative expression of *Efhd1* measured with qPCR in control and *Efhd1* overexpressed AML12 cells are shown in bar plot.

Abbreviations: Con, Control AML12 cells; Efhd1-OE, EFHD1 overexpressed AML12 cells
